# Theta-phase locking of single neurons during human spatial memory

**DOI:** 10.1101/2024.06.20.599841

**Authors:** Tim A. Guth, Armin Brandt, Peter C. Reinacher, Andreas Schulze-Bonhage, Joshua Jacobs, Lukas Kunz

## Abstract

The precise timing of single-neuron activity in relation to local field potentials may support various cognitive functions. Extensive research in rodents, along with some evidence in humans, suggests that single-neuron activity at specific phases of theta oscillations plays a crucial role in memory processes. Our fundamental understanding of such theta-phase locking in humans and its dependency on basic electrophysiological properties of the local field potential is still limited, however. Here, using single-neuron recordings in epilepsy patients performing a spatial memory task, we thus aimed at improving our understanding of factors modulating theta-phase locking in the human brain. Combining a generalized-phase approach for frequency-adaptive theta-phase estimation with time-resolved spectral parameterization, our results show that theta-phase locking is a strong and prevalent phenomenon across human medial temporal lobe regions, both during spatial memory encoding and retrieval. Neuronal theta-phase locking increased during periods of elevated theta power, when clear theta oscillations were present, and when aperiodic activity exhibited steeper slopes. Theta-phase locking was similarly strong during successful and unsuccessful memory, and most neurons activated at similar theta phases between encoding and retrieval. Some neurons changed their preferred theta phases between encoding and retrieval, in line with the idea that different memory processes are separated within the theta cycle. Together, these results help disentangle how different properties of local field potentials and memory states influence theta-phase locking of human single neurons. This contributes to a better understanding of how interactions between single neurons and local field potentials may support human spatial memory.

## Introduction

The activity of single neurons is embedded into waxing and waning local field potentials that represent the combined electrical activity of small neuronal populations within the neurons’ vicinity (*1–5*). A major component of local field potentials are fluctuations in the theta-frequency range at around 4–8 Hz (*6–12*). They are prominent in electrophysiological recordings from the rodent hippocampus and occur consistently when the animal navigates through its spatial environment (*7*, *13–17*). This “theta rhythm” may play a critical role in neural computation by grouping and segregating neuronal assemblies (*7*, *18*, *19*) and may thus be involved in cognitive functions including memory (*6*, *7*, *20–22*). In recent years, studies with epilepsy patients performing virtual and real-world navigation and memory tasks have described similar fluctuations of hippocampal local field potentials in humans, both in high and low theta-frequency ranges (1–5 and 6–10 Hz, respectively) (*8*, *16*, *23–30*). Human theta oscillations appear less clear and stable than in rodents, however (*8*, *9*, *16*, *30*). It thus remains unclear to what extent their putative properties and functions—including their interactions with single-neuron activity—generalize from rodents to humans.

A prominent observation on the relationship between single neurons and theta-frequency local field potentials is theta-phase locking (*31–36*). This phenomenon describes that a neuron activates at similar theta phases over time, for example at theta peaks or troughs. Theta-phase locking contrasts with other theta phase-dependent phenomena such as theta-phase precession where single neurons activate at successively earlier theta phases over time (*13*, *37–41*). In rodents, theta-phase locking has been described in a variety of neurons including spatially-modulated neurons representing locations and directions (*31*, *37*, *42–45*). Due to its high prevalence both within and across brain regions, theta-phase locking has been suggested to organize the activity of single neurons into neuronal assemblies (*22*, *30*, *36*), to enable interregional communication between neurons (*32*, *33*, *36*, *46*), and to trigger which neurons become linked during encoding to participate in the same replay events during consolidation (*42*). Phase locking of neuronal assemblies at distinct theta phases may furthermore help separate different mnemonic processes such as encoding and retrieval (*45*, *47*, *48*).

Based on this foundational work mostly performed in rodents, previous studies using single-neuron recordings in epilepsy patients started investigating the question whether theta-phase locking is present in the human brain as well, and whether it might be involved in human memory processes (*34*, *35*). These studies showed that, during a virtual spatial navigation task, neurons in widespread brain regions including hippocampus, amygdala, and parahippocampal regions were phase locked to oscillations in the theta-frequency range (*34*). Similar to observations in rodents (*32*, *33*), theta-phase locking also occurred between neurons in extrahippocampal regions and the hippocampal theta rhythm in patients performing another set of virtual spatial navigation tasks (*36*). In a recognition memory task with static images, stronger theta-phase locking was associated with successful memory formation as assessed with spike-field coherence (*35*). Similarly, in patients performing a verbal free recall task, a subset of neurons whose firing rates exhibited a subsequent memory effect showed stronger theta-phase locking during successful versus unsuccessful encoding events (*49*). This study also provided first evidence for theta-phase shifts between encoding and retrieval in human neurons (*49*). Together, these prior observations indicate that human single-neuron activity is related to local field potentials in the theta-frequency range and they suggest that these relationships support memory in humans.

Inspired by this previous work, we aimed at extending our basic understanding of theta-phase locking in the human medial temporal lobe in this study. Using human single-neuron recordings during a virtual navigation task with separate and self-paced periods for memory encoding and retrieval (*50–53*), we specifically aimed at investigating theta-phase locking as a function of different electrophysiological properties of the local field potential and across different memory states. Because human theta oscillations are relatively unstable over time (*24*, *25*, *27*, *30*) and considerably variable in frequency, we used a generalized-phase approach (*54*) to identify theta phases and employed time-resolved spectral parameterization (*55*, *56*) to distinguish between periods of strong versus weak periodic and aperiodic activity. Our results show that theta-phase locking is a prominent phenomenon in regions of the human medial temporal lobe and that it is strongly dependent on electrophysiological properties of the local field potential. Neuronal theta-phase locking was largely stable across different memory states and many neurons showed preferred theta phases that were similar between encoding and retrieval, with a few neurons showing significant shifts between encoding and retrieval. These findings provide new insights into the analysis and properties of human theta-phase locking and they may help identify their relevance for human memory processes.

## Results

### Spatial memory task

To investigate determinants of theta-phase locking in the human medial temporal lobe, we performed direct neurophysiological recordings in epilepsy patients (*57*, *58*) during a spatial memory task (*50*) (Table S1). In this “Treasure Hunt” task, participants navigated a virtual environment and were asked to encode and remember the locations of objects within the environment (Fig. 1; Methods). Briefly, during the encoding period of each trial, participants encountered two or three objects in treasure chests positioned at random locations on a virtual beach. They were then passively moved to an elevated recall position from where they could oversee the beach. After a short distractor period, participants performed cued recall. During location-cued object recall, they were cued with the locations and asked to speak aloud the names of the objects they had found at these locations. During object-cued location recall, they were presented with the objects and indicated the objects’ remembered locations on the beach. In a full session, participants aimed at encoding and remembering a total of 100 unique object–location associations.

**Figure 1.**
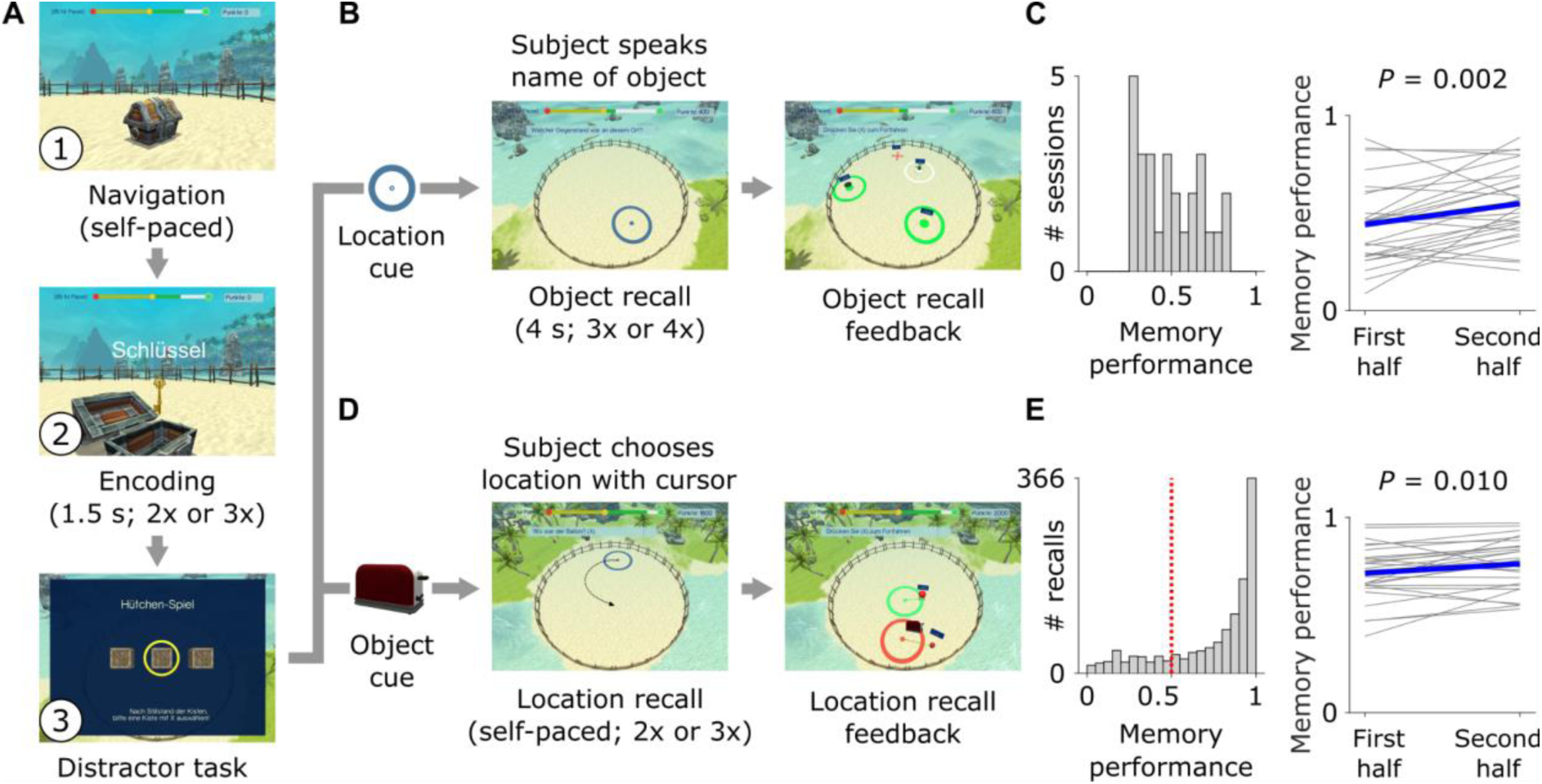
Spatial memory task. (**A**) During each navigation–encoding period of the task, participants freely navigated a virtual beach environment (1) and successively encountered two or three unique objects in treasure chests at random locations on the beach (2). Participants aimed at encoding the objects and their associated locations to recall them during the retrieval period. After the navigation–encoding period, participants were moved to an elevated recall position where they completed a distractor task (3) in order to prevent the participants from actively rehearsing the objects and their associated locations. After the distractor task, participants performed two types of memory recall. (**B**) During location-cued object recall, participants were presented with a location on the beach and recalled the associated object they had found at that location during encoding. (**C**) Memory performance during object recall per session (left) and improvement of object-recall memory performance over time, i.e., during the first half of all trials versus the second half of all trials (right). (**D**) During object-cued location recall, participants viewed an object and aimed at recalling the associated location on the beach. (**E**) Trial-wise memory performance during location recall (left) and improvement of location-recall memory performance over time (right). Red dotted line, chance performance. Gray lines, session-wise data; blue line, average.

Eighteen epilepsy patients participated in the task and contributed a total of 27 sessions. Participants performed the task for 60 ± 2 minutes (mean ± SEM), and their memory performance was similar to previous studies with this task (*50–52*) (Fig. 1, C and E). Participants showed memory-performance improvements over time for both types of memory recall, indicating that they successfully acquired knowledge of the spatial environment (object recall: paired *t*-test between the first and second half of all trials, *t*(26) = -3.402, *P* = 0.002; location recall: paired *t*-test, *t*(26) = -2.786, *P* = 0.010; *n* = 27 sessions; Fig. 1, C and E). Better performance during object recall was correlated with better performance during location recall (Spearman’s *rho* = 0.879, *P* < 0.001, *n* = 27 sessions). This task thus provided us with the opportunity to investigate theta-phase locking during memory encoding and retrieval, and as a function of memory performance.

### Human single neurons phase lock to the local theta rhythm

We performed direct neural recordings with high spatiotemporal resolution using Behnke-Fried microelectrodes (*57–61*) that allowed us to identify the activity of 1,025 neurons in various regions of the human medial temporal lobe (Fig. 2A; Fig. S1). We excluded neurons with fewer than 25 spikes (i.e., action potentials) in total or no spikes in more than 80% of segments across any trial condition, resulting in a total number of 666 neurons for all analyses.

**Figure 2.**
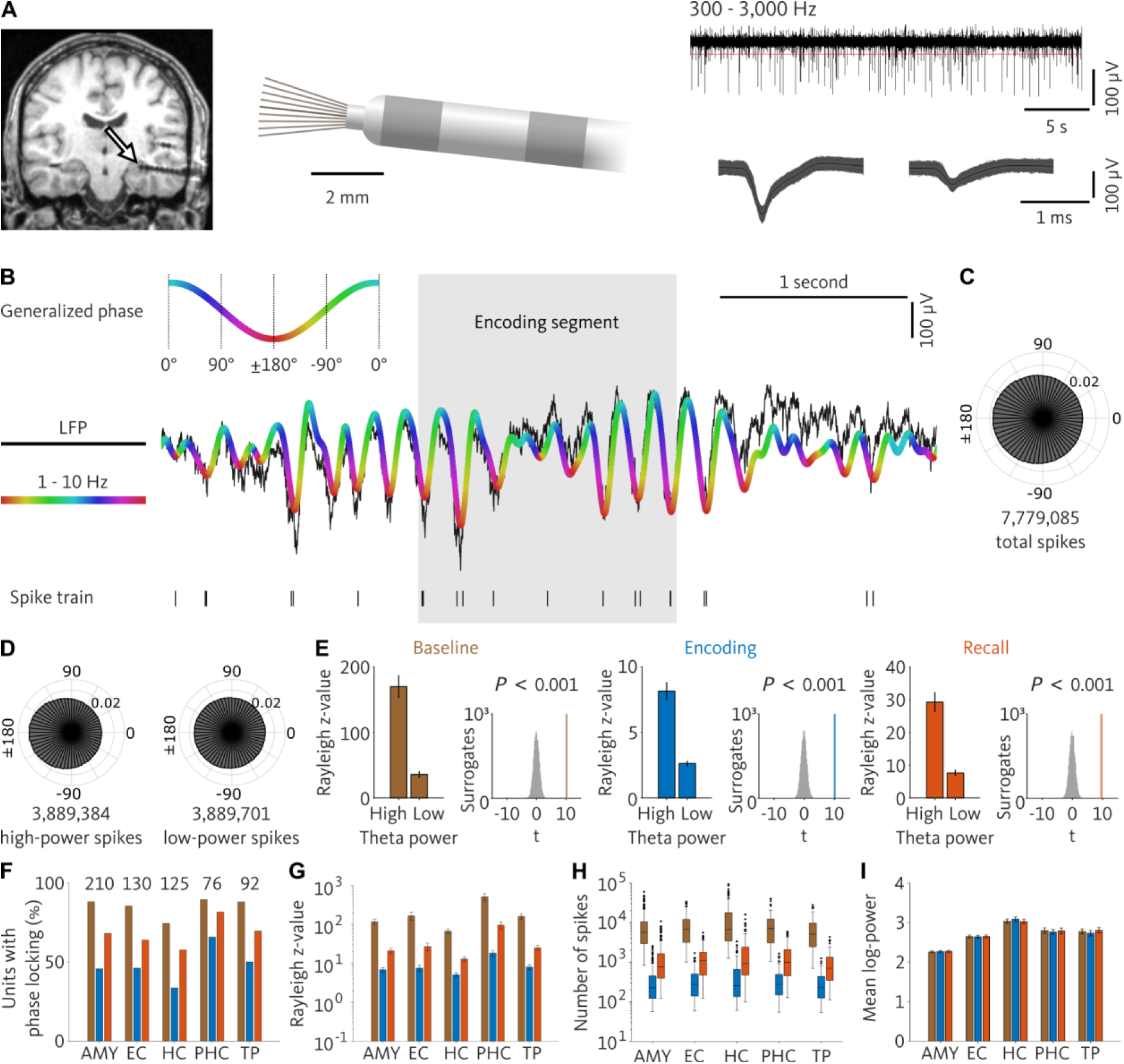
Neural recordings and general theta-phase locking. (**A**) Microelectrode recordings. Left, example post-implantation MRI. Arrow points at the tip of the Behnke–Fried electrode. Middle, schematic of the inner end of a Behnke– Fried electrode. Microelectrodes protrude from the tip of the macroelectrode by 3–5 mm. Top right, local field potential filtered between 300–3,000 Hz. Horizontal red line, threshold for spike detection. Bottom right, spike waveforms of two units recorded on this microelectrode. (**B**) Analysis procedure. We filtered the local field potential in a broad theta-frequency range (1–10 Hz) and computed the generalized phase of the filtered trace. For each spike, we extracted its corresponding theta phase. (**C**) General theta-phase locking across the entire experiment. Spikes pooled across all neurons preferentially occurred at the theta trough. (**D**) Theta-phase locking across the entire experiment as a function of whether the spikes coincided with high or low theta power (median split), pooled across all units. (**E**) Average theta-phase locking during high and low theta power across units, quantified with the Rayleigh *z*-value. Results are shown for baseline (left; entire session except for encoding and recall periods), encoding (middle) and recall (right). We compared the Rayleigh *z*-values between high and low power using *t*-tests and ranked the empirical *t*-values within surrogate *t*-values (plots to the right). *P*-values are Bonferroni corrected for three tests. Rayleigh *z*-values should not be compared between trial periods as they are strongly influenced by spike counts. (**F–I**) Theta-phase locking, number of spikes, and theta power in different brain regions during baseline (brown), encoding (blue) and recall (orange). (**F**) Percentages of units with significant phase locking. Numbers at top indicate the total number of neurons per region. (**G**) Rayleigh *z*-values across units. Error bars, standard error of the mean. (**H**) Number of spikes per region, across neurons. Box plots show medians, 25th and 75th percentiles, and outliers as dots. (**I**) Log-transformed power assigned to the spikes of each neuron. Error bars, standard error of the mean across neurons. AMY, amygdala; EC, entorhinal cortex; HC, hippocampus; PHC, parahippocampal cortex; TP, temporal pole.

We also extracted the low-frequency component of the microelectrode data in the theta-frequency range (1–10 Hz) using a generalized phase approach (*54*) to characterize the timing relationship between the neurons’ spiking activity and the theta rhythm. We opted for this approach with a broad filter band of 1–10 Hz as our visual inspection of the data identified many periods in which the local field potential showed fluctuations and oscillatory activity with rapidly changing frequencies. Our preliminary analyses with the filter-Hilbert method including narrow-band filtering often failed to fit these fluctuations neatly, and we thus decided to treat them as a coherent entity and opted for the generalized phase approach instead. This led to a good fit between the original and the filtered signal in the theta-frequency range (e.g., Fig. 2B; Fig. S2).

In a first step, we investigated general theta-phase locking of neuronal spikes across the entire task, without distinguishing between periods with or without clear theta oscillations, as done in prior studies (*34*). To quantify each neuron’s phase-locking strength, we calculated the Rayleigh *z*-value for the distribution of theta phases at which the neuron’s spikes occurred. To assess statistical significance, we ranked the empirical *z*-value within surrogate *z*-values that we obtained by circularly shifting the theta phases relative to the action potentials and recomputing the Rayleigh test (alpha level, 0.05; Methods). We observed many neurons for which the spikes were strongly locked to a particular phase of the local field potentials filtered in our broad theta-frequency range (571 of 666 neurons; 86%; binomial test versus 5% chance, *P* < 0.001). For example, a neuron in the left entorhinal cortex of a participant preferentially spiked at the theta troughs (surrogate analysis with circular shift of phase data, Rayleigh *z* = 5.089 × 10^3^, *P* < 0.001, *n* = 10,261 spikes; Fig. 2B). Pooling all spikes from all neurons, we found that they were generally clustered around the theta trough (Rayleigh test, *z* = 2.673 × 10^4^, *P* < 0.001, *n* = 7,779,085 spikes; Fig. 2C). This finding replicates previous observations of strong theta-phase locking in the human medial temporal lobe (*34*, *36*) and indicates that there is a tight temporal relationship between single-neuron activity and local low-frequency activity in the human medial temporal lobe.

### Theta-phase locking is increased for spikes associated with high theta power

Next, we aimed at identifying whether theta-phase locking is modulated by basic properties of the local field potential. We thus asked whether theta-phase locking was stronger during periods of elevated theta power. We found that theta-phase locking was present both during periods with high and during periods with low theta power using a median split (Rayleigh tests: high, *z* = 2.851 × 10^4^, *P*_corr._ < 0.001, *n* = 3,889,384 spikes; low, *z* = 3.918 × 10^3^, *P*_corr._ < 0.001, *n* = 3,889,701 spikes; Bonferroni corrected for two tests). When we directly compared theta-phase locking strengths between high versus low theta power, we found that theta-phase locking was more strongly expressed during periods with high theta power (surrogate analysis with spike label shuffling, Δ*z =* 2.459 × 10^4^, *P* < 0.001; Fig. 2D; Fig. S3A), which is in line with previous results (*34*). In designing the surrogate analysis, we ensured that the exact number of spikes in both conditions was maintained in all surrogate rounds, which is relevant as the Rayleigh *z*-value increases with the spike count (Fig. S4).

We also tested whether this modulation of theta-phase locking by theta power varied between different trial periods including encoding, recall, and baseline (where the baseline period comprised the entire session except for encoding and recall periods). In each of these three experimental periods, we separated the spikes of each unit into the groups of high and low theta power using a median split. We found that phase-locking was consistently stronger in the high-power than in the low-power condition for baseline, encoding, and recall periods (surrogate analysis with spike label shuffling: baseline, *t*(665) = 10.135, *P*_corr._ < 0.001; encoding, *t*(665) = 10.097, *P*_corr._ < 0.001; recall, *t*(665) = 9.608, *P*_corr._ < 0.001; Bonferroni corrected for three tests; Fig. 2E; Fig. S3A). This result shows that the modulation of theta-phase locking by theta power is a general phenomenon that is consistent across different memory states, at least in our spatial memory task.

### Phase locking to the local theta rhythm varies across medial temporal lobe regions

We next tested whether theta-phase locking differed between medial temporal lobe regions, including the amygdala, entorhinal cortex, hippocampus, parahippocampal cortex, and temporal pole. We found that the percentage of cells with significant phase locking varied between the different brain regions (χ^2^ tests: baseline, χ^2^(4) = 15.327, *P*_corr._ = 0.012; encoding, χ^2^(4) = 16.198, *P*_corr._ = 0.008; recall, χ^2^(4) = 13.729, *P*_corr._ = 0.025; *n* = 666; Bonferroni corrected for three tests; Fig. 2F). Numerically, the percentage of phase-locking neurons was lowest in the hippocampus and highest in the parahippocampal cortex. To better understand the origin of this finding, we analyzed the units’ Rayleigh *z*-values and found that they differed between brain regions as well (ANOVA for linear mixed-effects model: fixed effect of region, *F*(4) = 15.693, *P* < 0.001; Fig. 2G). Post-hoc pairwise comparisons showed that the regional differences were driven by increased theta-phase locking in the parahippocampal cortex (Table S2). As lower theta-phase locking is equivalent with a higher variability of spiking-related theta phases that may encode additional information, we speculate that this result points at an increased potential for theta-phase coding in the other, non-parahippocampal medial temporal lobe regions.

We aimed at ruling out that this finding was a side-effect of other properties, as stronger theta-phase locking can result from higher spike counts (Fig. S4) and increased theta power (Fig. 2E). We thus performed control analyses to test whether spike counts or theta power varied between medial temporal lobe regions. This showed that spike counts did not differ between medial temporal lobe regions (Kruskal–Wallis tests: baseline, *H*(4) = 8.135, *P*_corr._ = 0.260; encoding, *H*(4) = 4.308, *P*_corr._ = 1; recall, *H*(4) = 10.161, *P*_corr._ = 0.113; Bonferroni corrected for three tests; Fig. 2H). In contrast, spike-associated theta power varied between medial temporal lobe regions (ANOVA for linear mixed-effects model: fixed effect of region, *F*(4) = 204.063, *P* < 0.001; Fig. 2I). The direction of this effect could not explain the regional differences in phase-locking strength, however, as we observed highest theta power in the hippocampus where we observed the lowest number of phase-locking units (Fig. 2F; Table S3). These control analyses suggest that regional differences in theta-phase locking are not simply reducible to differences in spike counts or theta power.

### Theta-phase locking is stronger during periods with high aperiodic slopes

Local field potentials comprise aperiodic (non-oscillatory) and periodic (oscillatory) components (*55*), both of which may reflect and influence different types of neural processing (*62*). Both components can also lead to changes in absolute theta power (*6*, *55*) (Fig. S5) and may thus underlie our observation that higher theta-phase locking emerges during periods of elevated theta power (Fig. 2E). We thus aimed at understanding whether theta-phase locking was influenced by non-oscillatory and oscillatory activity.

To this end, we utilized Spectral Parameterization Resolved in Time (SPRiNT) that enabled us to disentangle aperiodic and periodic activity in a temporally resolved manner (*56*). Briefly, we estimated power spectra in short, 3-s time windows every 500 ms over the course of each session (Fig. 3A). For each power spectrum, we then calculated the slope of the aperiodic activity and identified whether at least one oscillatory peak in our broad theta-frequency range (1–10 Hz) was present or not (Fig. 3B). This approach allowed us to group neuronal spikes as a function of high versus low aperiodic slopes (median split) and according to whether they occurred in the presence or absence of theta oscillations (Fig. 3A).

**Figure 3.**
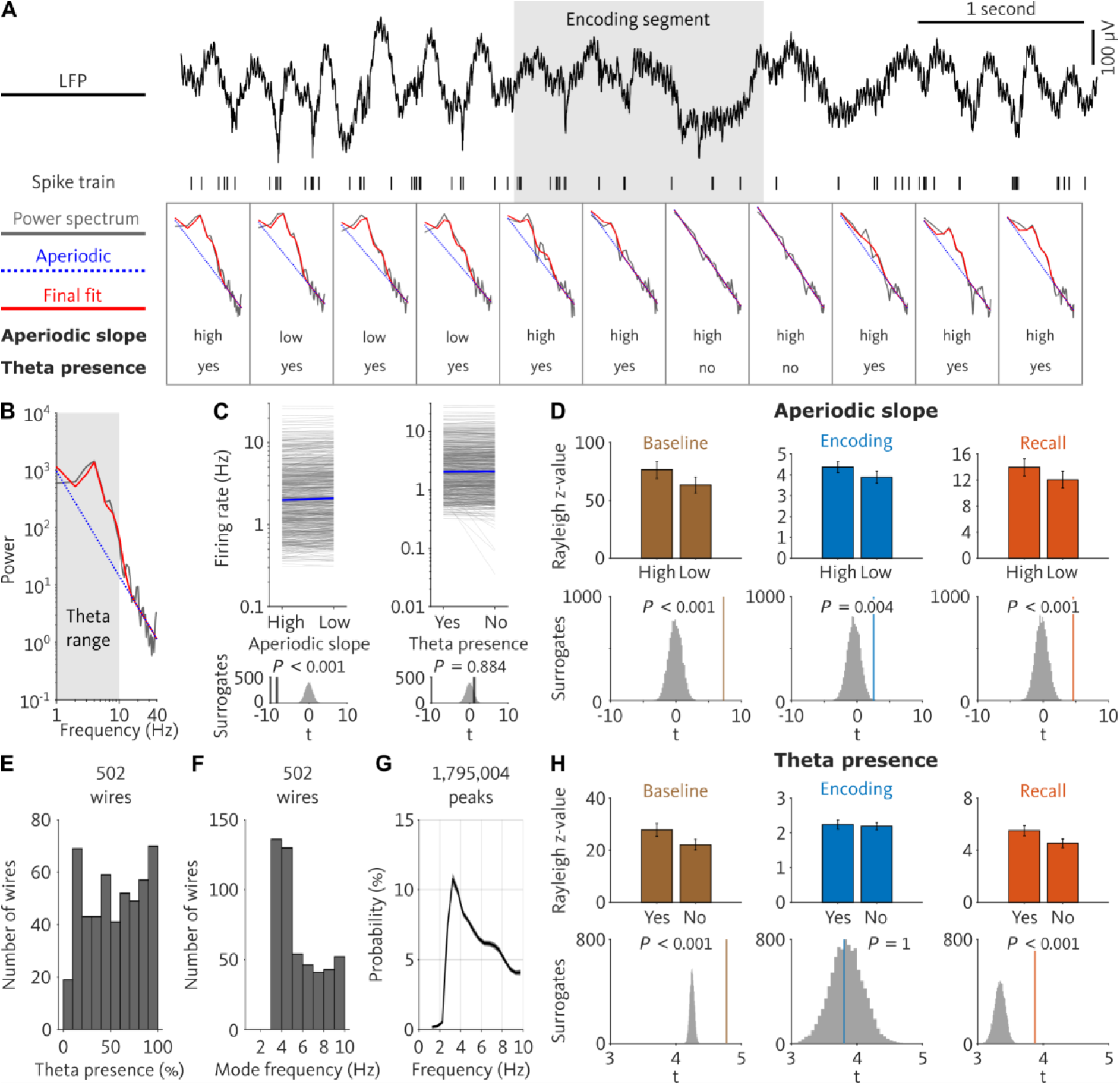
Phase locking as a function of aperiodic slope and the presence of theta oscillations. (**A**) We utilized Spectral Parameterization Resolved in Time (SPRiNT) (*56*) to analyze power spectra of the local field potentials in time steps of 500 ms. For each power spectrum (1–40 Hz; gray), SPRiNT provided a fit to its aperiodic component (dotted blue line) and a final fit to the original power spectrum (red). We grouped time windows into those with “high” versus “low” aperiodic slopes (median split). We also grouped them into time windows with versus without theta oscillations (“yes” and “no”, respectively) depending on whether SPRiNT detected a narrowband peak in the theta-frequency range. (**B**) Example power spectrum to illustrate aperiodic and final fit. Gray area, theta-frequency range. (**C**) Firing rate as a function of aperiodic slope (left) and as a function of theta-oscillation presence (right). Each line, one unit. Blue lines connect median firing rates. Bottom: Empirical *t*-values of paired *t*-tests comparing firing rates between conditions (black vertical lines), along with distributions of surrogate *t*-values (gray histograms). Surrogates were obtained by randomly shuffling the condition labels. (**D**) Theta-phase locking during high and low aperiodic slopes for baseline (left), encoding (middle), and recall (right). Empirical *t*-values of paired *t*-tests were ranked in surrogate distributions of *t*-values to assess significance (bottom). *P*-values are Bonferroni-corrected for three tests. (**E**) Histogram of percentages of time windows with theta oscillations. (**F**) Histogram of mode frequencies in the theta-frequency range (1–10 Hz). (**G**) Probability distribution of center frequencies of theta peaks (see Fig. S6 for results from different brain regions). Gray shaded area, standard error of the mean across microwires. (**H**) Theta-phase locking during the presence versus absence of theta oscillations, separately for baseline (left), encoding (middle), and recall (right). For visualization, Rayleigh *z*-values were computed based on subsampled data so that both conditions contained the same number of neuronal spikes (Fig. S4). Empirical *t*-values of paired *t*-tests were ranked in surrogate distributions of *t*-values to assess significance (bottom). *P*-values are Bonferroni-corrected for three tests.

In a preliminary step, we analyzed whether neuronal firing rates were modulated by aperiodic slopes and found that periods with high aperiodic slopes were associated with decreased neuronal firing rates (surrogate analysis with label swapping, *t*(665) = -8.337, *P* < 0.001; Fig. 3C; Fig. S3B). This observation is in line with the idea that higher (i.e., more negative) aperiodic slopes are related to increased inhibition (*62*) and thus lead to a suppression of neuronal activity. In contrast, firing rates during time windows with theta oscillations were not different from those without theta oscillations (surrogate analysis with label swapping, *t*(664) = 1.200, *P* = 0.884; Fig. 3C; Fig. S3B).

We then examined the impact of aperiodic slopes on neuronal theta-phase locking. We found that theta-phase locking was stronger for neuronal spikes associated with high aperiodic slopes as compared to those associated with low aperiodic slopes (median split). This effect was consistent across baseline, encoding, and recall periods (surrogate analysis with spike label shuffling: baseline, *t*(665) = 7.275, *P*_corr._ < 0.001; encoding, *t*(664) = 2.551, *P*_corr._ = 0.004; recall, *t*(665) = 4.557, *P*_corr._ < 0.001; Bonferroni corrected for three tests; Fig. 3D; Fig. S3A). Different firing rates during high versus low aperiodic slopes (see above) could not explain this result as our median split ensured the same number of spikes in both conditions. This finding indicates that the aperiodic slope of the local field potential is a major determinant of the strength of theta-phase locking.

### Phase locking is stronger during periods with theta oscillations

So far, we used all data to investigate theta-phase locking without excluding periods that did not show clear theta oscillations. This approach was driven by previous studies (*34*) and because we wanted to approach possible determinants of theta-phase locking as broadly as possible. We next asked how the presence of theta oscillations in the local field potentials modulated theta-phase locking and hypothesized that theta-phase locking would be stronger during periods with clear theta oscillations as compared to periods without them. Using SPRiNT, we thus identified all 3-s short time windows that contained at least one oscillatory peak within the theta-frequency range of 1–10 Hz.

In a preliminary step, we characterized the presence of theta oscillations in our recordings. For all 502 microwires, we computed the percentage of time windows with theta oscillations and found that theta oscillations were present about 54% of the time, with a high variability across microwires (Fig. 3E). To understand whether theta oscillations occurred at a preferred theta frequency, we computed the mode center frequency of all detected peaks between 1 and 10 Hz across wires (1,795,004 peaks in total) and observed that they occurred most often at 3–5 Hz (Fig. 3, F and G). Comparable frequency distributions of theta oscillations have been observed in prior studies in which participants performed similar virtual navigation tasks (*16*, *30*, *36*). Performing this analysis separately for microwires from different brain regions revealed distinct region-specific distributions of theta peak frequencies (Fig. S6). In the hippocampus, we observed a bimodal distribution with frequency peaks at 3 and 8 Hz. This is in line with the existence of both low and high theta in the human hippocampus (*26*, *49*, *63*).

We then analyzed how the presence of theta oscillations modulated phase locking during encoding, retrieval, and baseline periods. During baseline and recall (but not during encoding) we found that phase locking was stronger for neuronal spikes occurring during theta oscillations as compared to those outside clear theta oscillations (surrogate analysis with spike label shuffling: baseline, *t*(662) = 4.773, *P*_corr._ < 0.001; encoding, *t*(644) = 3.802, *P*_corr._ = 1; recall, *t*(645) = 3.876, *P*_corr._ < 0.001; Bonferroni corrected for three tests; Fig. 3H; Fig. S3A). The null effect during encoding may be due to reduced statistical power. Overall, both non-oscillatory and oscillatory components of the local field potential appear to modulate theta-phase locking in the human brain.

### Human single neurons theta-phase lock during memory encoding and retrieval

We were curious whether neuronal theta-phase locking varied as a function of memory state and performance. We thus investigated theta-phase locking separately for memory encoding and retrieval. Across all 666 neurons, we found 306 neurons that exhibited theta-phase locking during encoding and 444 neurons with theta-phase locking during retrieval (binomial tests versus 5% chance, both *P*_corr._ < 0.001, Bonferroni corrected for two tests). 273 units exhibited theta-phase locking during both encoding and retrieval (binomial test, *P* < 0.001; Fig. 4A). When only considering encoding and retrieval segments with successful memory performance, we found 163 neurons that exhibited phase locking during both encoding and retrieval, and 190 such neurons when only using segments with unsuccessful performance (binomial tests, both *P*_corr._ < 0.001, Bonferroni corrected). These results show that theta-phase locking is prevalent during both encoding and retrieval, and that the number of phase-locking neurons is similar during periods of successful and unsuccessful memory performance.

**Figure 4.**
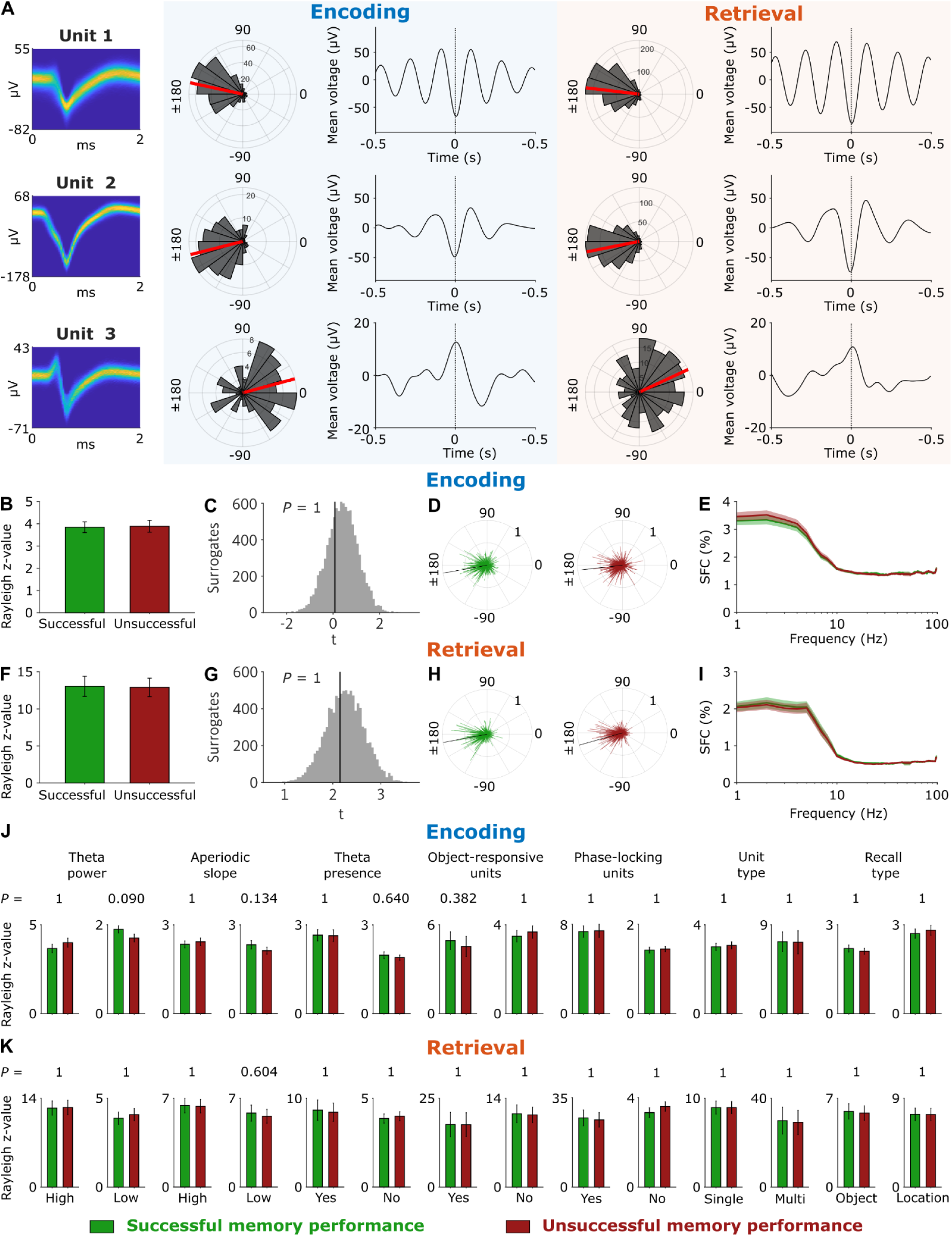
Theta-phase locking of human single neurons during memory encoding and retrieval and as a function of memory performance. (**A**) Examples of neuronal theta-phase locking during encoding and retrieval. Polar histograms show the theta phases of the units’ spikes during encoding (left) and retrieval (right). Red lines, mean phase angles. 1–10 Hz bandpass-filtered spike-triggered local field potentials are shown for encoding (left) and retrieval (right). Spike waveforms are shown as density plots. (**B**) Theta-phase locking during successful versus unsuccessful encoding. Error bars, standard error of the mean across units. (**C**) Empirical and surrogate *t*-values for comparing the Rayleigh *z*-values between successful and unsuccessful encoding. *P*-value is Bonferroni corrected for testing both encoding and retrieval. (**D**) Summary plots across all units showing their preferred theta phases and mean resultant vector lengths during successful (left) and unsuccessful (right) encoding. Black lines, mean phase across units. (**E**) Spike-field coherence for successful and unsuccessful encoding. Shaded areas, mean ± standard error of the mean across units. (**F**) Theta-phase locking during successful versus unsuccessful retrieval. Error bars, standard error of the mean across units. (**G**) Empirical and surrogate *t*-values of comparing the Rayleigh *z*-values from successful and unsuccessful retrieval. *P*-value is Bonferroni corrected for testing both encoding and retrieval. (**H**) Summary plots across all units showing their preferred theta phases and mean resultant vector lengths during successful (left) and unsuccessful (right) retrieval. Black line, mean phase across units. (**I**) Spike-field coherence for successful and unsuccessful retrieval. (**J**) Theta-phase locking during successful versus unsuccessful encoding for different data subsets. No significant differences were observed. Recall type refers to object-cued recall and location-cued recall, where the encoding process is the same but the recall processes differ. (**K**) Theta phase locking during successful versus unsuccessful retrieval for different data subsets. No significant differences were observed. SFC, spike-field coherence.

We next tested whether the strength of theta-phase locking was higher during segments with successful versus unsuccessful memory performance, separately for encoding and retrieval. We did not find stronger theta-phase locking (i.e., higher Rayleigh *z*-values) during successful encoding, indicating that the neurons’ preference toward a particular theta phase during encoding was independent of whether encoding was successful or not (surrogate analysis with spike label shuffling, *t*(665) = 0.078, *P*_corr._ = 1, Bonferroni corrected for two tests; Fig. 4, B and C; Fig. S3A). To corroborate this observation, we calculated the spike-field coherence in a frequency-resolved way between 1 and 100 Hz (*35*). We did not find any frequency with stronger phase locking during successful versus unsuccessful memory encoding (all *P*_corr._ = 1, Bonferroni corrected for 100 frequencies and two tests; Fig. 4E).

When investigating recall periods, we similarly found that neuronal theta-phase locking did not differ between segments with successful versus unsuccessful memory performance (surrogate analysis with spike label shuffling, *t*(665) = 2.149, *P*_corr._ = 1, Bonferroni corrected for two tests; Fig. 4, F and G; Fig. S3A). A more general spike-field coherence analysis, as described above, did not reveal any frequencies between 1–100 Hz with stronger phase locking during successful versus unsuccessful retrieval (all *P*_corr._ = 1, Bonferroni corrected; Fig. 4I). In contrast to previous studies with recognition and verbal free recall memory tasks that observed stronger phase locking during successful memory encoding (*35*, *49*), these results suggest that the strength of theta-phase locking during memory encoding and retrieval did not differ as a function of memory success in our spatial memory task.

We additionally tested whether the neurons’ preferred theta phases changed between successful and unsuccessful memory. This analysis showed no significant differences in preferred theta phases between segments with successful versus unsuccessful memory performance (surrogate analysis with label swapping; encoding: *F*(665) = 0.652, *P*_corr._ = 0.718; retrieval: *F*(665) = 1.352, *P*_corr._ = 0.162, Bonferroni corrected for two tests; Fig. 4, D and H).

To provide a more comprehensive assessment, we then asked whether the relationship between phase locking and memory performance might be modulated by theta power, aperiodic activity, or the presence of theta oscillations. We also distinguished between units that responded with a firing-rate increase during encoding (“object-responsive units”; Fig. S7) and units that did not show such object responsiveness. We separately examined units with and without significant phase locking, multi units and single units, and unit activity during object recall or location recall. In all these cases, we found that neuronal theta-phase locking was similar between successful and unsuccessful segments, both for encoding and retrieval (surrogate analysis with spike label shuffling; all *P*_corr._ > 0.090, Bonferroni corrected for performing this analysis for encoding and retrieval and for two data conditions in each case; Fig. 4, J and K). Additional control analyses showed that time-frequency resolved power, which can modulate theta-phase locking effects, did not differ as a function of memory performance during encoding and retrieval (Fig. S8). In summary, our findings suggest that theta-phase locking strength and the preferred theta phases of single neurons during encoding and recall did not depend on whether memory retrieval was successful or not.

### Some neurons shift their preferred theta phases between encoding and retrieval

We considered that neurons might exhibit theta-phase locking to different theta phases during encoding versus retrieval (Fig. 5). Such theta-phase shifts have been predicted by the “Separate Phases of Encoding And Retrieval” (SPEAR) model and may help to avoid interference between encoding and retrieval processes (*47–49*, *64*). We thus tested for neurons that exhibited a significant phase difference between encoding and retrieval (Fig. 5A) and found evidence for such phase shifts in some neurons of our dataset (62 of 666 neurons; 9%; binomial test versus 5% chance, *P* < 0.001; Fig. 5, B, C and F). When only considering neurons that showed significant phase locking during both encoding and retrieval, we found a similar proportion of neurons with significant phase shifts between encoding and retrieval (26 of 273; 10%; binomial test, *P* < 0.001; Fig. 5, C and F).

**Figure 5.**
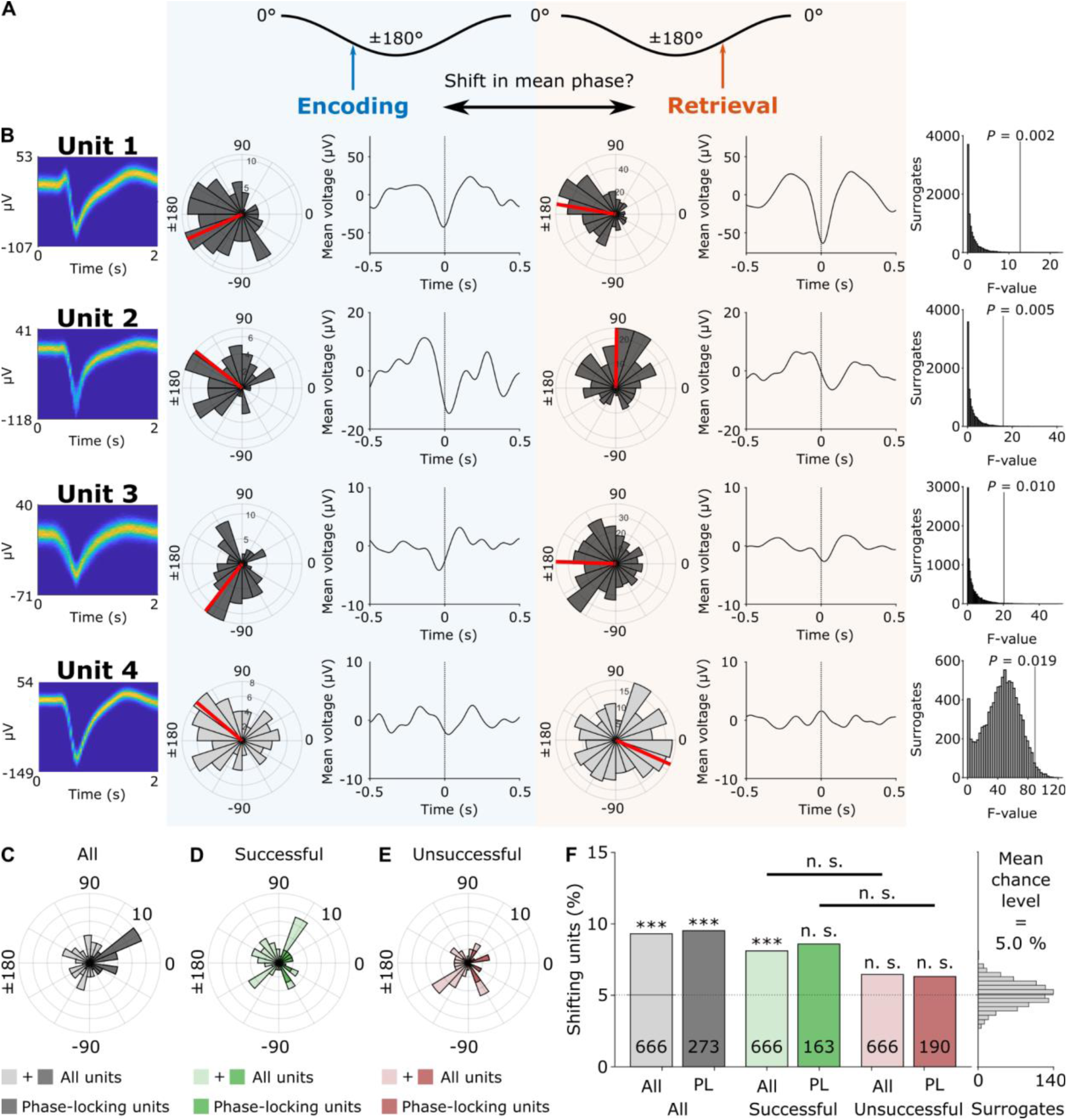
Shifts of preferred theta phases between encoding and retrieval. (**A**) Hypothesis that neurons shift their preferred theta phases between encoding and retrieval (*47*). (**B**) Four example units with significantly shifted theta phases between encoding and retrieval. Polar histograms show the theta-phase distributions of all spikes during encoding (blue) and retrieval (orange). Theta-filtered spike-triggered averages are shown next to the polar histograms. Left, spike-waveforms as density plots. Right, distributions of surrogate *F*-values (histogram) and empirical *F*-values (vertical lines) of Watson-Williams tests between the theta phases during encoding and retrieval. Units 1–3 significantly phase-locked during encoding and retrieval, and unit 4 did not. (**C–E**) Polar histograms show the angular differences between theta phases during encoding versus retrieval for neurons with significant theta-phase shifts between encoding and retrieval. Results are shown for all units and for units with significant theta-phase locking. (**C**) Results for all segments (*n* = 62 significant units from the pool of all units*; n* = 26 significant units from the pool of significantly phase-locking units). (**D**) Results for successful segments (*n* = 54 and *n* = 14). (**E**) Results for unsuccessful segments (*n* = 43 and *n* = 12). An angular difference of 0° corresponds to identical theta phases during encoding and retrieval. (**F**) Percentages of units with significant theta-phase shifts for all (transparent color) and significantly phase locking units (opaque color). Percentages are compared to 5% chance level. Gray histogram on the right, empirically estimated chance level using surrogate data, confirming the *a priori* chosen 5% chance level. \*\*\**P* < 0.001; n.s., not significant.

We then tested whether theta-phase shifts depended on memory performance. When only considering data from successful encoding and retrieval segments, we observed a significant number of neurons with significant phase shifts between encoding and retrieval (54 phase-shifting neurons of 666 neurons; 8%; binomial test, *P*_corr._ < 0.001, Bonferroni corrected for performing the test for successful and unsuccessful memory performance; Fig. 5, D and F). We observed a similar trend when only considering units with significant phase locking during both encoding and retrieval (14 phase-shifting neurons of 163 neurons; 9%; binomial test, *P*_corr._ = 0.070, Bonferroni corrected; Fig. 5, D and F). When only considering data from unsuccessful segments, the number of significantly shifting units was neither significant for all units (43 of 666; 6%, binomial test, *P*_corr._ = 0.110, Bonferroni corrected; Fig. 5, E and F) nor for units with significant phase locking during encoding and retrieval (12 of 190; 6%; binomial test, *P*_corr._ = 0.488, Bonferroni corrected; Fig. 5, E and F). These results suggest that phase shifts were slightly more prevalent during successful as compared to unsuccessful memory, though the direct comparisons were not significant (surrogate analysis with unit label shuffling: all units, Δ = 2%, *P* = 0.136; phase-locking units, Δ = 2%, *P* = 0.276).

Next, we quantified the absolute phase difference of phase-shifting units between encoding and retrieval and found that it was 85° ± 47° (mean ± circular SD). Among the neurons with significant phase locking during both encoding and retrieval, the absolute phase difference between encoding and retrieval was 36° ± 22° on average (mean ± SD; Fig. 5C). These effects were similar for successful and unsuccessful memory performance (successful, all neurons: 101° ± 45°; successful, phase-locking neurons: 42° ± 18°; unsuccessful, all: 109° ± 45°; unsuccessful, phase-locking neurons: 47° ± 32°; Fig. 5, D and E). The extent of these phase shifts was thus lower than originally proposed (*47*), but they may still be sufficiently large to help separate encoding and retrieval processes.

We finally asked whether theta-phase shifts varied as a function of other properties of the local field potential. We thus estimated the number of units with significant phase shifts separately for spikes associated with high and low theta power, high and low aperiodic slope, and for those during theta oscillations (Fig. S9). We found a significant number of phase-shifting units for high theta power (64 of 666; 10%; binomial test, *P*_corr._ < 0.001, Bonferroni corrected for two tests) but not for low theta power (37 of 666; 6%; binomial test, *P*_corr._ = 0.557, Bonferroni corrected; Fig. S9A). The number of phase-shifting neurons was also significant for spikes associated with high aperiodic slopes (51 of 651; 8%; binomial test, *P*_corr._ = 0.003, Bonferroni corrected) but not for those with low aperiodic slopes (41 of 652; 6%; binomial test, *P*_corr._ = 0.163, Bonferroni corrected; Fig. S9B). Similarly, the number of phase-shifting neurons was significant for spikes in the presence of theta oscillations (45 of 609; 7.4%; binomial test, *P*_corr._ = 0.013, Bonferroni corrected) but not in their absence (36 of 552; 6.5%; binomial test, *P*_corr._ = 0.131, Bonferroni corrected; Fig. S9C).

These results indicate that shifts in the preferred theta phase are—at least to some extent—modulated by electrophysiological properties including theta power, aperiodic slope, and the presence of theta oscillations. Overall, our findings of significant phase shifts between encoding and retrieval provide additional, albeit limited, evidence for the SPEAR model (*47*, *49*, *64*, *65*). As the majority of neurons showed similar theta phases between encoding and retrieval, however, we suggest that both theta-phase shifts and stable theta phases may contribute to encoding–retrieval processes.

## Discussion

In this study, we aimed at improving our basic understanding of neuronal theta-phase locking in humans. Our results demonstrate strong phase locking of single neurons to the local theta rhythm in different regions of the human medial temporal lobe. Several electrophysiological properties of the local field potentials including theta power, the presence of clear theta oscillations, and the slope of aperiodic activity modulated the strength of theta-phase locking. This indicates that theta-phase locking is a dynamic phenomenon that is influenced by a variety of properties of the local field potential. Furthermore, theta-phase locking was similarly pronounced during periods with successful versus unsuccessful memory performance, suggesting that additional studies are needed to pinpoint the memory relevance of theta-phase locking. While most neurons preferred similar theta phases between different memory states, some neurons showed significant shifts in their preferred theta phases between encoding and retrieval, roughly in line with the idea that different memory processes happen at distinct parts of the theta cycle (*47*). Together, these results help identify the electrophysiological determinants and functional properties of theta-phase locking in humans. This contributes to a better understanding of the relationship between single neurons and neuronal populations in the human brain.

### Determinants of theta-phase locking

When we estimated theta-phase locking across the entire task, we found a high number of neurons that exhibited theta-phase locking (about 86%; Fig. 2). Confining the analysis to the shorter periods of encoding and retrieval clearly reduced the number of theta-phase locking neurons. This indicates that the amount of data (and, thus, statistical power) is a major factor in determining the number of phase-locking units.

To compute theta phases, we used a generalized phase approach (*54*) including broad-band filtering between 1–10 Hz. This is different from previous phase-locking studies that estimated theta phases at various individual theta frequencies and computed phase locking for each of these frequencies separately (*34*, *35*, *49*). We opted for the generalized phase approach as visual inspection of our local field potentials showed that they were not stable and sustained narrowband oscillations, but often dynamic in frequency and amplitude over time. We observed that the generalized phase estimation fit the raw data best and led to fewer phase distortions. A prior study in monkeys showed that neuronal spiking is coupled more strongly to generalized phase than to narrowband theta or alpha phases (*54*). The use of generalized phase may thus constitute another reason for the high number of neurons with theta-phase locking in our study.

Single-neuron spiking occurred most often at the trough of the theta rhythm, similar to previous observations in humans (*34*). This effect was present both at the level of single neurons and when we pooled the spike phases of all neurons. This finding is in line with the view that theta peaks are associated with increased inhibition that prevents single neurons from spiking (*1*, *7*, *62*), imposing a major restriction on when action potentials can occur during the theta cycle. In this way, theta phases may gate and coordinate the output of single neurons (*19*, *65–69*).

We found that different signal properties of the local field potential were associated with theta-phase locking. First, we observed that theta-phase locking was more strongly expressed during periods with high theta power, again in line with prior findings (*34*). Higher theta power may thus reflect an additional layer of inhibition, further restricting the occurrence of single-neuron activity to parts of the theta cycle close to its trough. As it is a prominent observation in rodents that theta power increases with running speed (*63*, *70*) at which the animal samples sensory information at a higher rate, we speculate that tighter theta-phase locking during periods of increased theta power may be useful for processing rapidly incoming information in a precise and efficient manner.

Furthermore, using spectral parameterization in a time-resolved manner (*55*, *56*), we found stronger phase locking when aperiodic activity showed steeper (i.e., more negative) slopes and when periodic activity featured clear theta oscillations in the local field potential (Fig. 3). This again fits with the idea that higher levels of inhibition lead to more restricted single-neuron firing—and thus stronger phase-locking—as steeper slopes of aperiodic activity are considered markers of increased inhibition (*62*). Because recent studies have shown that aging and mental disorders affect theta oscillations and characteristics of aperiodic activity (*71–73*), we propose that neuronal theta-phase locking may change under these conditions as well.

Together, these results provide new insights into the electrophysiological and behavioral determinants of theta-phase locking in the human medial temporal lobe. They suggest that studies investigating theta-phase locking as a function of cognitive processes should consider changes in the basic properties of the local field potential as well. This may help delineate whether changes in theta-phase locking are a more or less independent substrate of cognitive functioning.

### Theta-phase locking during successful versus unsuccessful memory

Previous studies observed stronger theta-phase locking related to successful memory performance in human single-neuron recordings (*35*, *49*). These studies showed stronger theta-phase locking at frequencies of 1–9 Hz during successful memory formation in a recognition memory task with static images (*35*). They also showed stronger theta-phase locking in subsequent-memory effect neurons (those neurons with increased firing rates during successful encoding) at frequencies of 2–5 Hz during successful encoding in a verbal free recall task with words (*49*).

Here, we did not find stronger theta-phase locking during successful memory performance, neither during encoding nor during retrieval. We tested for these memory effects in several subgroups of the data, for example only including spikes in the presence of clear oscillations or only including object-responsive cells, and we did not find differences in theta-phase locking as a function of memory performance in any of these groups (Fig. 4). These findings differ from the above-mentioned studies that described a positive relationship between the strength of theta-phase locking and the successful encoding of images and words (*35*, *49*).

We speculate that the dynamic nature of our spatial memory task is relevant to explaining our null effect. Specifically, in contrast to studies with static images or words that have a clear-cut onset and are shown after a blank screen—which may lead to a phase reset of theta oscillations time-locked to stimulus onset (*20*)—our spatial memory task featured encoding and retrieval periods that were smoothly embedded into the other trial periods. As phase resets have been shown to be associated with successful encoding of static stimuli (*20*, *74*), we propose that stronger phase resets of theta oscillations—in combination with a consistent delay of single-neuron activity relative to stimulus onset—may underlie stronger theta-phase locking during successful memory encoding. Future studies may investigate this idea by testing whether theta-phase locking exhibits a memory effect for static but not for dynamic stimuli that can be explained through phase resets and stimulus-locked single-neuron activity.

### Theta-phase shifts between encoding and retrieval

Single-neuron activity occurring at different theta phases during encoding versus retrieval may be beneficial for memory by preventing interference between the learning of new information and the retrieval of old information (*47*, *48*). This idea has been articulated by the “separate phases of encoding and retrieval” (SPEAR) model (*47*). Specifically, this model suggests that encoding happens at the theta peak of local field potentials in the hippocampal CA1 region, when input from the entorhinal cortex is strong and input from CA3 is weak. Conversely, the model proposes that retrieval occurs at the theta trough of local field potentials in CA1, when input from the entorhinal cortex is weak and input from CA3 is strong (*48*).

Guided by this model, we investigated whether neurons in various regions of the human medial temporal lobe showed theta-phase locking at different theta phases during the encoding and retrieval periods of our spatial memory task. While we found that the majority of neurons locked to similar theta phases during encoding and retrieval, we also identified a small but significant number of neurons that shifted their preferred theta phases between encoding and retrieval (Fig. 5). Interestingly, these shifts were numerically slightly more pronounced during successful as compared to unsuccessful memory. Our results thus provide limited additional evidence for the SPEAR model, in line with earlier investigations of theta-phase shifts in rodents and humans (*49*, *64*).

Theta-phase shifts varied in their magnitude and direction between neurons. In neurons that showed significant theta-phase locking both during encoding and retrieval, the extent of theta-phase shifts was about 40° on average, and it was about 90° for neurons without significant theta-phase locking during encoding and retrieval. A previous human single-neuron study found theta-phase shifts of comparable magnitude (about 100°) during a verbal free-recall task when investigating neurons whose firing rates showed a subsequent memory effect (*49*). Similarly, in rodents, hippocampal CA1 neurons showed theta-phase shifts of less than 90° when rats navigated through novel versus familiar environments that were used to trigger encoding and retrieval states, respectively (*64*). These shifts are smaller in size than predicted by the SPEAR model, but they may still help to separate encoding and retrieval processes. Future studies may clarify the extent of theta-phase shifts, their distribution across different brain regions, and their prevalence during different types of memory.

### Limitations of this study

To examine theta-phase locking as broadly as possible, we analyzed theta-phase locking across the entire task and tested subsequently whether theta-phase locking was more strongly pronounced during periods with clear theta oscillations. It currently remains an open question whether it is useful to investigate theta-phase locking during periods without clear theta oscillations, and future studies on theta-phase locking may thus focus on periods with theta oscillations *a priori*. As human theta oscillations are variable in strength and frequency, however, the detection of theta oscillations and other types of oscillatory activity is non-trivial. In this study, we used time-resolved spectral parameterization (*56*) to identify periods with theta oscillations. Future studies may compare the performance of this procedure with other methods for detecting oscillatory activity, such as BOSC (*75*) and MODAL (*22*), to identify its advantages and disadvantages.

In addition to identifying periods with theta oscillations, time-resolved spectral parameterization also allowed us to estimate the slope of aperiodic activity over short time windows. This showed that theta-phase locking dynamically changed as a function of aperiodic activity, with stronger theta-phase locking during periods with steeper aperiodic slopes. We note, however, that time-resolved spectral parameterization involves a number of analysis settings due to the complexity of the analysis, and we observed some variability in how well power spectra were fit depending on particular settings. Specifically, spectral analysis at low frequencies is difficult as these frequencies are close to the lower bound of the power spectrum and the frequency of slow-theta peaks can thus be misidentified. Visual inspection of our data nevertheless showed that our final choice of settings led to a good fit of the original power spectra overall. Computational studies may help to pinpoint the optimal settings for time-resolved spectral parameterization of local field potentials in humans.

Another limitation of this study is the finding that the strength of theta-phase locking was not associated with memory performance. As discussed above, we currently believe that this null effect is due to the dynamic nature of our spatial memory task, which may prevent theta-phase resets at the onset of the encoding and retrieval periods. Relatedly, we did not find strong differences in theta-phase shifts between successful versus unsuccessful memory performance, perhaps because of statistical power. Additional studies are thus needed to better understand the role of stable and shifting theta phases for memory success in humans (*6*, *7*, *48*).

## Acknowledgements

We are very grateful to all patients who participated in this study. We thank the clinical team of the Freiburg Epilepsy Center, Freiburg im Breisgau, Germany, for their continuous support. We thank the members of the Jacobs Lab at Columbia University and the members of the Mormann Lab at the University Hospital Bonn for feedback. T.A.G., A.B., A.S.-B., and L.K. were supported by the Federal Ministry of Education and Research (BMBF; 01GQ1705A) and by NIH/NINDS grant U01 NS113198-01. T.A.G. was supported by a stipend of the German National Academic Foundation (Studienstiftung des deutschen Volkes). P.C.R. received research support from the Fraunhofer Society (Munich, Germany) and personal fees, travel support, and honoraria from Boston Scientific, Brainlab, and Inomed. J.J. was supported by NIH grant MH104606 and the National Science Foundation (NSF). L.K. received funding from the German Research Foundation (DFG; Projektnummer 447634521) and was supported by the return program of the Ministry of Culture and Science of the State of North Rhine-Westphalia.

## Author contributions

T.A.G. and L.K. developed hypotheses and designed the study; L.K., A.S.-B., and J.J. acquired funding; L.K., A.S.-B., P.C.R., and A.B. recruited study participants; P.C.R. implanted electrodes; L.K., A.B., and T.A.G. collected the data; T.A.G. analyzed the data; T.A.G., J.J., and L.K. discussed the results; T.A.G. and L.K. wrote the paper; all authors reviewed and revised the final manuscript.

## Competing interests

The authors declare no competing interests.

## Methods

### Human participants

We tested *N* = 18 human participants (10 female; age range, 22–63 years; mean age ± SEM, 41 ± 3 years), who were epilepsy patients undergoing treatment for pharmacologically intractable epilepsy at the Freiburg Epilepsy Center, Freiburg im Breisgau, Germany (Table S1). Written informed consent was obtained from all patients. The study conformed to the guidelines of the ethics committee of the University Hospital Freiburg, Freiburg im Breisgau, Germany.

### Spatial memory task

During experimental sessions, patients performed a computerized spatial memory task (“Treasure Hunt”), while seated in a hospital bed. The task was developed with Unity3D (Unity Technologies, San Francisco, CA, USA) and has been used and described in previous studies (*50–52*). The virtual environment of the task comprised a plain beach surrounded by a circular wooden fence. Outside the environment, the background scenery comprised multiple mountains, palm trees, barrels, the sea, and the vast sky. Participants performed up to 40 experimental trials. At the beginning of each trial, participants were placed at a random location on the virtual beach. Participants then navigated to successively appearing treasure chests (Fig. 1A), using a game controller. Upon reaching a chest, the chest opened to reveal an object and its name (Fig. 1A). After 1.5 seconds, the chest and object disappeared. During each navigation–encoding period, participants traveled to 2 or 3 chests. In a complete session, participants traveled to a total of 100 chests. After encountering the last chest of a trial, participants were moved to an overhead perspective of the environment. Participants then played a distractor game (Fig. 1A), in which they had to track the position of a coin hidden under one of three constantly moving boxes. The distractor game had a mean duration of 6.6 s.

After the distractor game, the retrieval period began. In each trial, participants completed either location-cued object recall or object-cued location recall. During object recall, *n* + 1 locations were successively shown on the beach (in random order), where *n* corresponds to the number of treasure chests encountered during the preceding navigation–encoding period. Participants had four seconds to speak out loud the name of the object that was contained in the treasure chest at the highlighted location or ‘‘Nichts’’ (German for ‘‘nothing’’) for the one location not associated with a treasure chest (Fig. 1B, left). Correctness of the response was evaluated using Cortana (Microsoft, Redmond, WA, USA) and manually checked (and corrected, if necessary) during data analysis. During object-cued location recall, the names of the objects contained in the *n* treasure chests from the preceding navigation–encoding period were shown to the participants in random order. Participants were asked to move a small target circle across the beach to the remembered location of the associated treasure chest (self-paced duration; Fig. 1D, left).

After being probed for all the locations/objects from a given trial, participants completed a recency-judgment task in which they were asked to judge which of two objects they had encountered later during the preceding navigation–encoding period (not analyzed in this study). Finally, participants received feedback on whether they had correctly recalled the object names (Fig. 1B, right) or locations (Fig. 1D, right) and how they performed in the recency-judgment task. The next trial started by transporting the participant back onto the beach.

To quantify the participants’ memory performance, we calculated two different metrics: object recall was evaluated based on whether a location-cued object was correctly recalled or not. Location-recall performance was quantified by calculating the Euclidean distance between the remembered location and the correct location of the cued object (‘‘drop error’’). Following previous studies (*50*, *52*, *76*), drop errors were ranked within one million potential drop errors to obtain normalized performance values. A value of 1 indicated the best possible response, while a value of 0 indicated the worst. We defined a location recall as successful, if its normalized performance value was higher than the session’s median normalized performance value.

We defined the electrophysiological data during an encoding or retrieval event as an “encoding segment” and as a “retrieval segment,” respectively. An encoding–retrieval pair was classified as “successful” if the participant correctly recalled the encoded item or its location during the retrieval phase. Any data from the paradigm that did not fall into the categories of encoding or retrieval segments were classified as “baseline segments.”

### Neurophysiological recordings

Patients underwent surgical implantation of intracranial depth electrodes in the medial temporal lobe to identify the epileptic seizure focus for potential subsequent surgical resection. Electrode numbers and locations varied across patients and were solely determined by clinical needs. Neuronal signals were recorded using Behnke–Fried depth electrodes (Ad-Tech Medical Instrument Corp., Racine, WI, USA). Each depth electrode contained a bundle of nine platinum–iridium microelectrodes with a diameter of 40 μm that protruded from the tip of the electrode (*57*). We recorded spikes and local field potentials from eight microelectrodes, with the ninth microelectrode serving as a reference. Microelectrode coverage included amygdala, entorhinal cortex, fusiform gyrus, hippocampus, insula, parahippocampal cortex, temporal pole, and visual cortex. We recorded the microelectrode data at 30 kHz using NeuroPort (Blackrock Microsystems, Salt Lake City, UT, USA). We aligned neurophysiological recordings and behavioral data using triggers that were sent from the paradigm to the recording system.

### Spike detection and sorting

Neuronal spikes were detected and sorted using Wave_Clus 3 (*77*). We used default settings with the following exceptions: ‘‘template_sdnum’’ was set to 1.5 to assign unsorted spikes to clusters in a more conservative manner; ‘‘min_clus’’ was set to 60 and ‘‘max_clus’’ was set to 10 in order to avoid over-clustering; and ‘‘mintemp’’ was set to 0.05 to avoid under-clustering. We evaluated all clusters visually based on the spike shape and its variance, inter-spike interval (ISI) distribution, and the presence of a plausible refractory period. If necessary, we manually adjusted or excluded clusters. Spike waveforms are shown as density plots in all figures (except for Fig. 2A).

In total, we identified *N* = 1,025 clusters on 649 wires across 27 experimental sessions from all 18 participants. Clusters from different sessions were treated as statistically independent units. An experienced rater (L.K.) assigned the tips of depth electrodes to brain regions based on post-implantation MRI scans in native space so that units recorded from the corresponding microelectrodes could be assigned to these regions. To assess recording quality (Fig. S1), we calculated the number of units recorded on each wire; the ISI refractoriness for each unit; the mean firing rate for each unit; and the waveform peak signal-to-noise ratio (SNR) for each unit. The ISI refractoriness was assessed as the percentage of ISIs with a duration of <3 ms. The waveform peak SNR was determined as: SNR = A_peak_/SD_noise_, where A_peak_ is the absolute amplitude of the peak of the mean waveform, and SD_noise_ is the standard deviation of the raw data trace (filtered between 300 and 3,000 Hz).

We classified a unit as a single unit based on the following criteria (*78*, *79*): (1) Visual inspection of spike shape and its variance; (2) ratio between peak of the mean waveform (i.e., spike amplitude) and standard deviation at the first sampling point (i.e., noise) greater than 3; and (3) <1% of the spikes with an inter-spike interval <3 ms, which takes into account that neurons have a refractory period. Units that did not fulfill the criteria of a single unit were considered as multi units.

To ensure sufficient statistical power, we excluded units with a low number of spikes. Specifically, we excluded units with fewer than 25 spikes during any of the following conditions: successful encoding, unsuccessful encoding, successful retrieval, or unsuccessful retrieval. To mitigate the likelihood that effects were driven by a few segments only, units with no spikes in more than 80% of the encoding or retrieval segments of any of these conditions were also excluded from all analyses. Of all 1,025 units, a total of 666 units on 502 wires met the inclusion criteria and were thus included in our analyses. We recorded *n* = 210 units from the amygdala, *n* = 130 units from the entorhinal cortex, *n* = 24 units from the fusiform gyrus, *n* = 125 units from the hippocampus, *n* = 2 units from the insula, *n* = 76 units from the parahippocampal cortex, *n* = 92 units from the temporal pole, and *n* = 7 from the visual cortex.

### Preprocessing of local field potentials

We preprocessed the local field potentials in several ways to prepare them for analyses of spike-field relationships. To minimize the contribution of spike shapes toward phase and power estimations, we computed the mean waveform of each unit in the 100–3000 Hz bandpass-filtered signal and subtracted it from the original local field potential whenever a spike occurred. The start and end samples of the mean spike were tapered to zero to prevent edge artifacts. We then resampled the local field potential to 2 kHz and applied a 4th order finite impulse response band-stop filter to remove electrical line noise at 48–52 Hz and its harmonics at 98–102 and 148–152 Hz. Finally, we demeaned the local field potential to center it around zero.

### Estimation of theta phase and theta power in the local field potential

To estimate theta phase and theta power, we filtered the preprocessed local field potentials between 1 and 10 Hz (comprising both the low- and high-theta frequency range) using an FIR filter (order = 6000). To extract instantaneous power and phase angles of our filtered signal, we used a single-sided Fourier transform and the previously established generalized phase approach (*54*). This approach enhances the accuracy of phase estimations in local field potentials filtered across a broad frequency range. It corrects for phase distortions introduced by low- and high-frequency intrusions, ensuring that the phase information reflects the true broadband fluctuations of the local field potential. We chose a 1 Hz cut-off for low-frequency intrusions. For a detailed explanation of the generalized phase approach and how it corrects for low- and high-frequency intrusions see the original publications (*54*, *80*). For each spike, we then identified the temporally closest phase and instantaneous power (Fig. 2B). Since we filtered in the 1–10 Hz frequency range we refer to the assigned values as “theta phase” and “theta power.”

### Detection of interictal epileptiform discharges

We automatically detected interictal epileptiform discharges (IEDs) in the preprocessed local field potentials following the same methods and detection criteria as described before (*76*). Timepoints with high amplitudes, sharp amplitude changes and power increases across a broad range of frequencies were considered belonging to an IED and excluded from further analyses. Spikes occurring during these timepoints were excluded from all analyses. We do not refer to IEDs as interictal spikes, because we reserved the term ‘spike’ for extracellularly recorded single-unit action potentials throughout the paper.

### Theta-phase locking

To investigate if there was consistent phase-locking towards a specific phase of the local field potential we visualized the distribution of phases assigned to the spikes of all units in a polar histogram (Fig. 2C). For power-specific analyses, we used a median split of the spikes of each unit, categorizing them as either “high” or “low” based on their theta power (Fig. 2D). We performed Rayleigh tests to quantify the non-uniformity of the resulting theta-phase distributions. The Rayleigh *z-*statistic was derived using the equation *z* = *n* × *r*², where *n* is the number of spikes, and *r* is the mean resultant vector length (MRVL). The MRVL quantifies the consistency of phase angles. A value close to 0 indicates low phase locking and a value close to 1 indicates high phase locking.

To assess the strength of theta-phase locking and the preferred theta phase for each unit, we calculated the Rayleigh *z*-value and the circular mean of the theta phases at which the spikes occurred. For each unit, we ranked the empirical *z*-value within 1,001 surrogate *z*-values, which we obtained by circularly shifting the theta-phase data relative to the spike time series (Fig. S3C). A unit was considered to exhibit significant theta-phase locking, if the empirical *z*-value exceeded the 95^th^ percentile of all surrogate *z*-values.

### Theta-phase locking as a function of theta power

To compare phase locking between spikes with high and low power, we calculated the empirical difference of the two Rayleigh *z*-values and ranked it in a distribution of 10,001 *z*-value differences obtained by randomly shuffling the spike labels “high” or “low” (Fig. S3A). As the Rayleigh *z*-value increases with the number of spikes (Fig. S4), we ensured that the exact number of spikes in both conditions was maintained in all surrogate rounds.

We examined the modulation of theta-phase locking by theta power across different experimental periods: encoding, recall, and baseline (where the baseline period comprised the entire session except for encoding and recall periods). For each period, we separated neuronal spikes into high and low theta power groups based on a median split. We then computed Rayleigh *z*-values for both high and low power spikes of each unit, followed by a paired *t*-test to compare these *z*-values across units (Fig. 2E). We ranked the empirical *t*-value among 10,001 surrogate *t*-values (Fig. S3A). The surrogates were generated by randomly assigning each spike to either the “high” or “low” power condition, while preserving the number of spikes in each condition. As the Rayleigh *z*-value increases with the number of spikes, a direct comparison of *z*–values between baseline, encoding, and retrieval is not possible, due to the different number of spikes in these conditions.

### Theta-phase locking as a function of brain region

We investigated whether phase locking varied across different brain regions during encoding, retrieval, and baseline. We included only brain regions with data from at least five sessions: amygdala, entorhinal cortex, hippocampus, parahippocampal cortex, and temporal pole. We first compared the number of significantly phase-locking units with the number of units with no significant phase locking across brain regions using Chi-squared tests (Fig. 2F). Resulting *P*-values were Bonferroni corrected for applying these tests on the three experimental periods baseline, encoding, and retrieval. We next compared Rayleigh *z*-values across units using a linear mixed-effects model (dependent variable: Rayleigh *z*-values; fixed effect: brain region; random effects: experimental session and period; Fig. 2G).

We also examined regional differences in spike counts and spike-associated theta power, given their potential effects on phase locking. Due to the heterogeneous distribution of spike counts, we decided to use non-parametric Kruskal–Wallis tests to compare spike counts across brain regions (Fig. 2H). We applied three tests, each corresponding to one experimental period, and Bonferroni-corrected the resulting *P*-values. To compare spike-associated theta power (Fig. 2I), we computed the average log-transformed theta power assigned to the spikes of each unit and compared the results using a linear mixed-effects model (dependent variable: log-transformed theta power; fixed effect: brain region; random effects: experimental session and period).

### Spectral Parameterization Resolved in Time

In addition to periodic, narrowband oscillations, local field potentials typically display a characteristic power distribution known as 1/f-like aperiodic activity, which appears as a straight line when plotted in log-log space. The slope of this aperiodic activity might reflect the balance of neuronal excitation and inhibition with steeper slopes reflecting stronger inhibition (*62*).

We utilized the previously established Spectral Parameterization Resolved in Time (SPRiNT) approach to disentangle aperiodic and periodic activity in a temporally resolved manner (*56*). SPRiNT leverages spectral parameterization (*55*) on short time windows. Our analyses showed that the outcome of the SPRiNT algorithm exhibited variable results based on different configuration settings. We thus adhered to default settings unless our visual inspection identified specific configurations that enhanced the algorithm’s compatibility with our dataset.

We applied SPRiNT to the 2 kHz resampled local field potentials of each session. The algorithm first computed short-time Fourier transforms from 1-second time windows with 50% overlap. Every 500 ms, it then averaged the power spectra from five consecutive time windows. Power spectra containing automatically detected interictal epileptiform discharges or artifacts were excluded from further analyses. For spectral parameterization, the algorithm fitted an aperiodic component to each average power spectrum within the 1 to 40 Hz frequency range. To distinguish periodic activity from the aperiodic component, up to three Gaussian peaks were identified. Peaks required a signal-to-noise ratio >2 and a height >0.5. We set the maximum peak width to 6 Hz (default = 12 Hz) to extract only narrow oscillatory peaks. We set the minimum peak width to 2 Hz (default = 0.5 Hz) after observing artificial peaks at stereotypical frequencies that lacked clear correlates in the power spectra. This adjustment effectively reduced the presence of these artifacts. We applied a proximity threshold to remove a peak if its center frequency fell within one standard deviation (default, 2 SD) of a relatively larger peak’s center frequency or within one standard deviation (default, 2 SD) from the edge of the frequency window. We adjusted this threshold to facilitate fitting peaks closer to the spectrum’s edge. The SPRiNT algorithm includes a post-processing step to eliminate transient outlier peaks, often caused by erroneously modeled noise. Using this procedure, we removed peaks that were not part of a cluster with at least three similar peaks (defined by a maximum frequency deviation <2.5 Hz) within six adjacent time bins.

### Firing rate as a function of aperiodic slope and theta oscillations

As a control analysis, we assessed the influence of aperiodic slope and theta oscillations on neuronal firing rates. To this end, we computed the aperiodic slope for each 500-ms time window using SPRiNT. We then split the time windows into “high” slope time windows (indicating a steeper negative slope) and “low” slope time windows (indicating a flatter negative slope) using a median split. We calculated the overall firing rate during high and low slope periods for each neuron. We used a paired *t*-test to compare firing rates between high and low slope conditions. To assess significance, we ranked the empirical *t*-value within a distribution of *t*-values derived from 10,001 surrogate datasets. These surrogate datasets were generated by randomly swapping the labels “high slope” and “low slope” for each neuron’s firing rates with a 50% probability. To investigate the influence of theta oscillations on neuronal firing rates, we performed the same analysis with the only difference being that we split the data into two groups based on the presence or absence of theta oscillations in each 500 ms time window.

### Phase locking as a function of aperiodic slope

We aimed at understanding the impact of aperiodic slope on neuronal theta-phase locking. To this end, we assigned each spike the aperiodic slope value derived from its corresponding 500 ms SPRiNT time window. We then did a median split of the spikes of each unit, categorizing them as either “high” or “low” based on their slopes. Similar to the power-specific analyses, we computed Rayleigh *z*-values for both high and low slope spikes of each unit, followed by a paired *t*-test to compare the *z*-values between high and low slope across units. To assess significance, we ranked the empirical *t*-value within 10,001 surrogates (Fig. S3A). These surrogates were generated by randomly assigning each spike to either the “high” or “low” slope condition. We performed this analysis separately for spikes during baseline, encoding, and retrieval.

### Analysis of theta oscillations

We next analyzed the occurrence of theta oscillations in the local field potentials. We identified all 500 ms SPRiNT time windows containing at least one peak with a center frequency within the theta range of 1 to 10 Hz. We then computed the percentage of time windows with theta oscillations for all 502 wires (Fig. 3E). We rounded the center frequencies of all detected peaks to the nearest integer and computed the mode frequency between 1 and 10 Hz for each wire (Fig. 3F). We also calculated the probability distribution of center frequencies for all theta peaks on each wire, using a 0.5 Hz frequency resolution (Fig. 3G; Fig. S6).

### Phase locking as a function of theta oscillations

We analyzed the impact of the presence of theta oscillations on theta-phase locking for spikes during baseline, encoding, and retrieval. To this end, we determined whether a spike occurred within a 500-ms SPRiNT time window where theta oscillations were detected or not. We computed Rayleigh *z*-values for spike phases in the presence versus absence of theta oscillations, followed by a paired *t*-test to compare the resulting *z*-values across units.

During data analysis, we noticed that the Rayleigh *z*-values were strongly modulated by the number of spikes per condition (Fig. S4). To address this issue, we implemented the following two steps: First, we ranked the resulting *t*-value of the paired *t*-test among 10,001 surrogates. These surrogates were generated by randomly reassigning each spike to either the “theta” or “no theta” condition, while maintaining equal spike counts for both conditions in each surrogate dataset (Fig. S3A). This approach ensured that any potential bias introduced by varying spike numbers was preserved in the surrogates. Second, for clearer visualization of our results, we performed a subsampling procedure on our original data. For each neuron, we identified the condition (“theta” or “no theta”) with the higher spike count. We then randomly drew 10,001 subsamples from the larger condition, ensuring that each subsample matched the spike count of the smaller condition. We calculated the mean Rayleigh *z*-value across these 10,001 subsamples, and used these *z*-values for plotting.

### Theta-phase locking during successful and unsuccessful encoding and retrieval

To assess and compare theta-phase locking strength and theta-phase angles at the unit level between segments associated with successful versus unsuccessful memory performance, we performed the following procedure separately for encoding and retrieval. We extracted all spike phases during successful and unsuccessful segments and computed Rayleigh *z*-values and the mean phases of the two groups. We then performed a paired *t*-test to compare the *z*-values of the units’ spike phases, and we performed a Watson–Williams test to compare the preferred theta phases during successful versus unsuccessful segments.

Next, we ranked the empirical *t*-value of the *t*-test in a distribution of 10,001 surrogate *t*-values and the empirical *F*-value of the Watson–Williams test in a distribution of 10,001 surrogate *F*-values. Surrogates were obtained by randomly reassigning each spike to either the “successful” or “unsuccessful” condition, while maintaining equal spike counts for both conditions in each surrogate dataset (Fig. S3A). This approach ensured that any potential bias introduced by varying spike numbers was preserved in the surrogates. The units’ theta-phase locking strength was considered significantly higher for successful versus unsuccessful memory performance if the empirical *t*-value was above 95% of the surrogate *t*-values. We considered the units’ preferred theta phases to differ between successful and unsuccessful memory performance, if the empirical *F*-value of the Watson-Williams test was above 95% of the surrogate *F*-values. The resulting *P*-values were Bonferroni corrected to account for the fact that we performed the test for both encoding and recall segments.

For clearer visualization of our results, we performed a subsampling procedure on our original data. For each neuron, we identified the condition (“successful” or “unsuccessful”) with a higher spike count. We then randomly drew 10,001 subsamples from the larger condition, ensuring that each subsample matched the spike count of the smaller condition. We calculated the mean Rayleigh *z*-value across these 10,001 subsamples, and used these *z*-values for plotting.

### Spike-field coherence

We used spike-field coherence (SFC) to quantify phase locking during successful and unsuccessful memory performance for both encoding and retrieval segments (*35*, *81*) in a frequency-resolved way (Fig. 4E). SFC measures the percentage of the power spectrum of the spike-triggered average (STA) over the average power spectrum of the local field potentials around the spikes, which were used to compute the STA. For each frequency, the SFC takes values between 0% and 100%. High SFC values indicate strong phase locking at a specific frequency (*35*).

To compute the SFC, we first extracted 500 ms of the 2 kHz resampled and mean-spike-removed local field potentials before and after each spike. Next, we resampled the local field potentials to a sampling rate of 250 Hz. The spike-triggered average was obtained by averaging the resulting traces. The power spectrum of the STA and the power spectra of the spike-triggered local field potential traces were calculated at linearly spaced frequencies from 1 to 100 Hz with intervals of 1 Hz using the FieldTrip implementation for multitaper analysis (*82*, *83*). We used Slepian tapers and a 4-Hz width of frequency smoothing. When comparing the SFC between two conditions, we used the same number of spikes for each condition. We randomly selected the number of spikes of the condition with fewer spikes for the condition with more spikes. This subsampling procedure was repeated 100 times and the resulting 100 SFC values were averaged for each frequency. To test for significant differences between successful and unsuccessful segments, we performed a paired *t*-test across units at each frequency bin and Bonferroni-corrected the *P*-values for 100 comparisons (*35*) and for performing the analysis separately for encoding and retrieval.

### Memory effects on phase locking for different subgroups

To investigate memory-related effects possibly restricted to specific parts of our dataset, we divided it into subgroups to compare phase-locking strength between successful versus unsuccessful memory performance. In each subgroup analysis, we applied a Bonferroni correction for four comparisons given that we tested two data groups (e.g., high and low theta power) and two trial periods (encoding and retrieval).

We grouped the data into the following subgroups: (1) separation of spikes based on a median split of their associated theta-power values; (2) separation of spikes based on a median split of their aperiodic slopes; (3) separation of spikes based on the presence versus absence of theta oscillations; (4) separation of units based on whether they were object-responsive or object-non-responsive units; (5) separation of units based on whether they exhibited theta-phase locking; (6) separation of units based on whether they were single units or multi units; and (7) separation of recall periods based on whether they were object-recall or location-recall periods.

We identified object-responsive units as those units that exhibited a significant increase in firing during the encoding period of the task. To this end, we calculated firing rates for each encoding segment and its corresponding baseline segment, the latter defined as the 1.5 seconds preceding the encoding period. We then compared the firing rates between encoding and baseline segments using a paired *t*-test. We ranked the resulting *t*-value within a distribution of 10,001 surrogate *t*-values that we obtained by randomly swapping/keeping the labels “encoding” and “baseline” for each encoding–baseline pair (Fig. S3B). Units with a *t*-value greater than 95% of the surrogate *t*-values were considered object-responsive units.

### Theta-phase shifts between encoding and retrieval

We also performed analyses to investigate how many units exhibited a significant phase shift between encoding and retrieval. For each unit, we used a Watson–Williams test to compare the phase distribution of all its spikes during encoding segments with the phase distribution of all its spikes during retrieval segments. We then created 10,001 surrogate datasets by randomly shuffling the segment labels “encoding” and “retrieval” (Fig. S3D) and performed a Watson–Williams test for every surrogate dataset. We ranked the empirical *F*-value of the unit’s Watson–Williams test in the distribution of the 10,001 surrogate *F*-values and considered a unit to exhibit a significant theta-phase shift between encoding and retrieval if the empirical *F*-value exceeded the 95^th^ percentile of all surrogate *F*-values.

To confirm that the 95^th^ percentile was chosen correctly when evaluating the statistical significance of units, we performed the following control analysis: We concatenated all encoding and retrieval phases in the order of their occurrence and circularly shifted the resulting list of phases by a random lag. We then reassigned the same number of spikes and their theta phases from the shifted list to the corresponding encoding and retrieval segments. This way, we created 1,001 surrogate datasets on which we performed the same analysis as the one we had performed on the empirical data. By calculating the average percentage of shifting units in each of the surrogate datasets, we obtained a distribution whose mean confirmed our *a priori* chosen alpha level of 5% (Fig. 5F).

We performed the analysis of phase-shifting neurons separately for all units and phase-locking units, and we separately analyzed all, successful, and unsuccessful segments (Fig. 5, C–F). We evaluated whether the number of shifting units observed in the empirical data was higher than the chance level of 5%. We also tested for significant phase shifts as a function of high versus low theta power, high versus low aperiodic slope, and the presence versus absence of theta oscillations.

### Code and data availability

Data analyses were performed with custom Matlab code and using the Matlab toolboxes CircStat (*84*) and FieldTrip version 20210912 (*83*). Custom Matlab code generated during this study for data analysis is available at https://github.com/NeuroGuth/GuthPhaseLocking2024. Additional information and data will be provided by the corresponding authors upon reasonable request.

---

**Figure S1.**
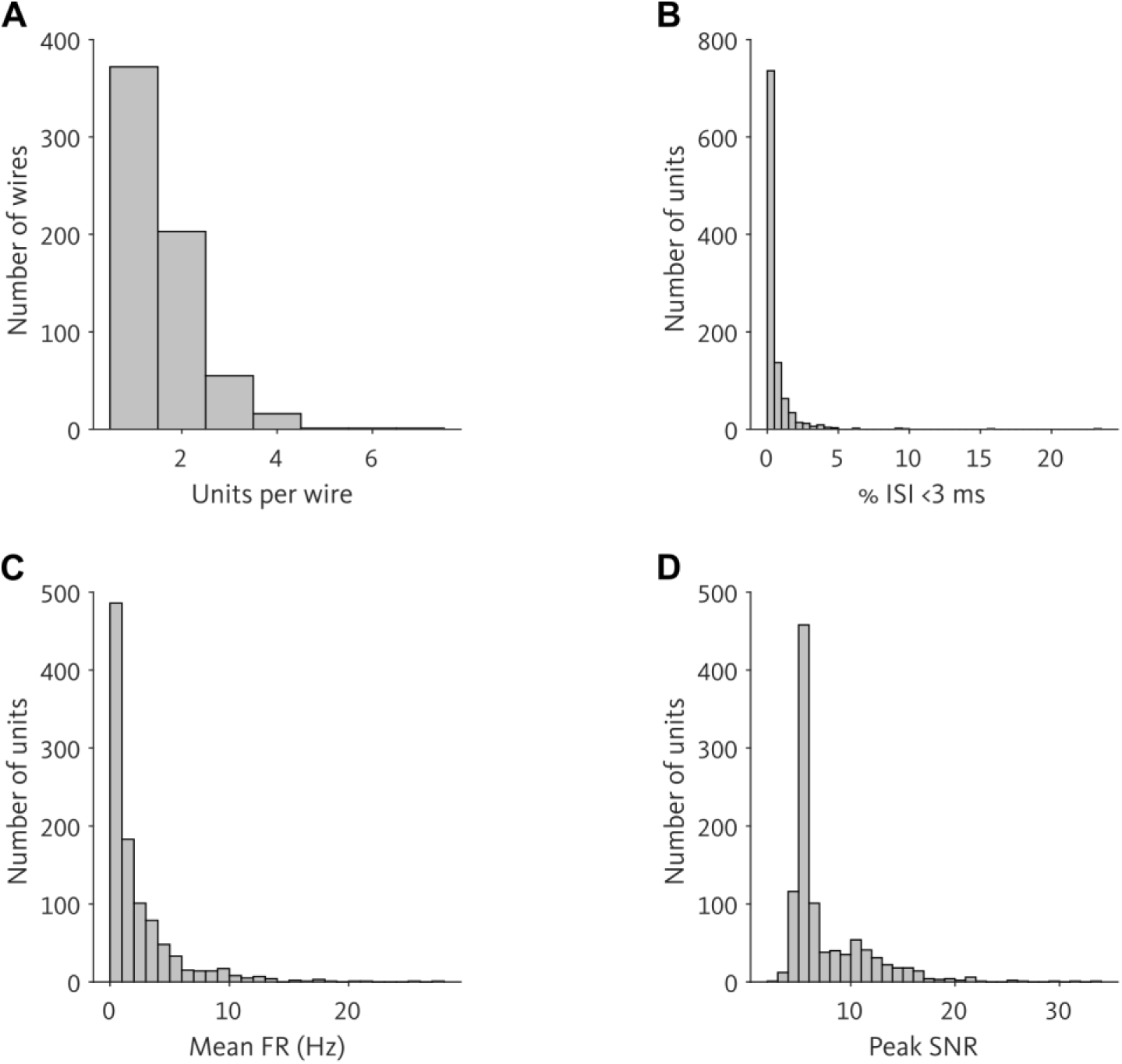
Quality assessment of single-neuron recordings. (**A**) Histogram of units per wire. On average, 1.58 ± 0.03 (mean ± SEM) units per wire were recorded. (**B**) Histogram of the percentages of inter-spike intervals that were shorter than 3 ms. On average, units exhibited 0.54 ± 0.04% (mean ± SEM) inter-spike intervals that were shorter than 3 ms. There were 7 units with values >5%. (**C**) Histogram of mean firing rates. On average, units exhibited mean firing rates of 2.31 ± 0.10 Hz (mean ± SEM). (**D**) Histogram of the mean waveform peak signal-to-noise ratio of each unit. On average, the signal-to-noise ratio of the mean waveform peak was 7.56 ± 0.12 (mean ± SEM).

**Figure S2.**
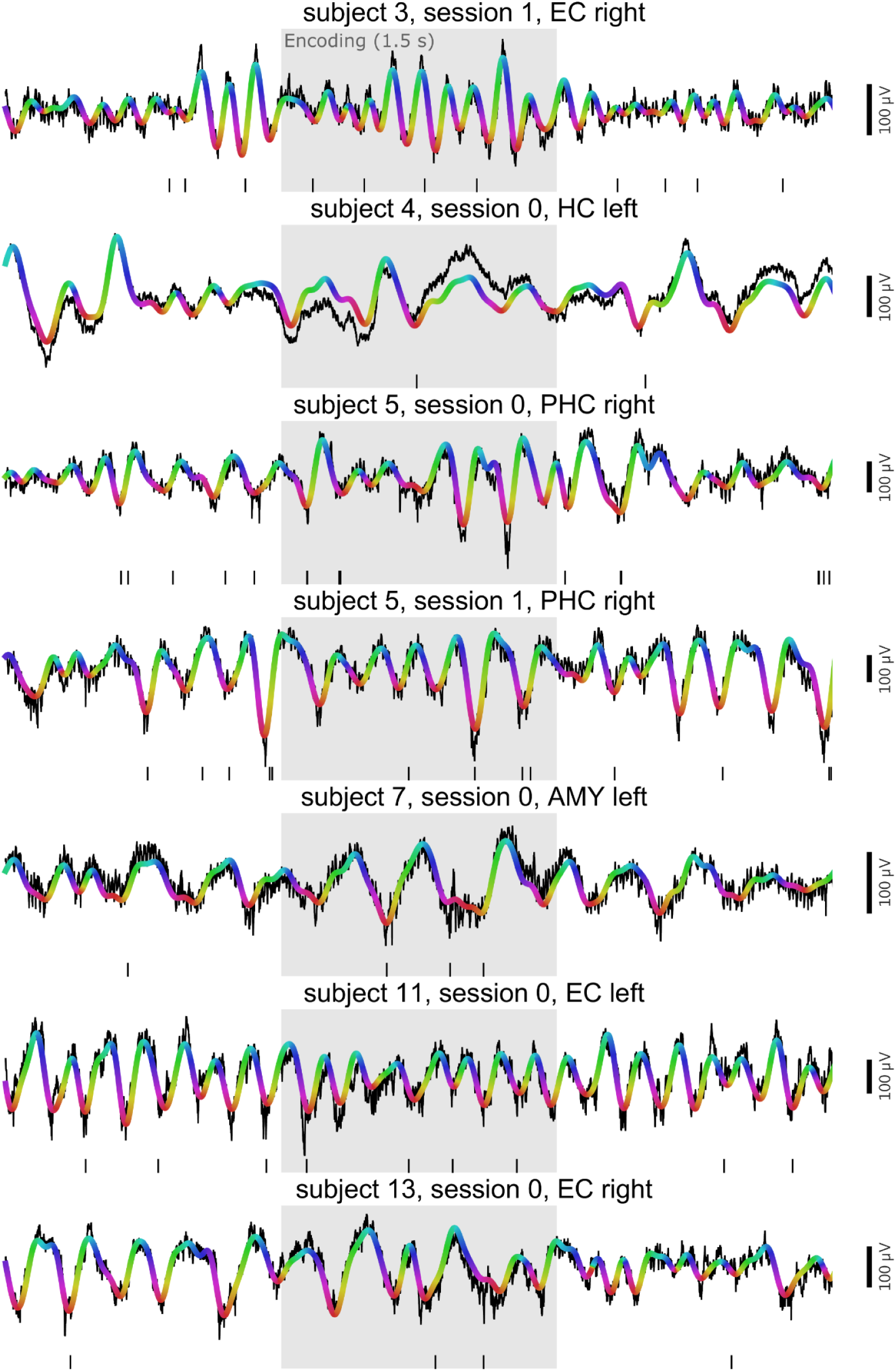
Examples of neuronal theta-phase locking in the human medial temporal lobe. Examples of raw (black) and filtered (1–10 Hz; colored) local field potentials from different experimental sessions. Spike trains are shown as vertical lines below the local field potentials. Color indicates the instantaneous theta-phase angle estimated using a generalized phase approach (as in Fig. 2B), and gray areas indicate the 1.5-s encoding period. AMY, amygdala; EC, entorhinal cortex; HC, hippocampus; PHC, parahippocampal cortex.

**Figure S3.**
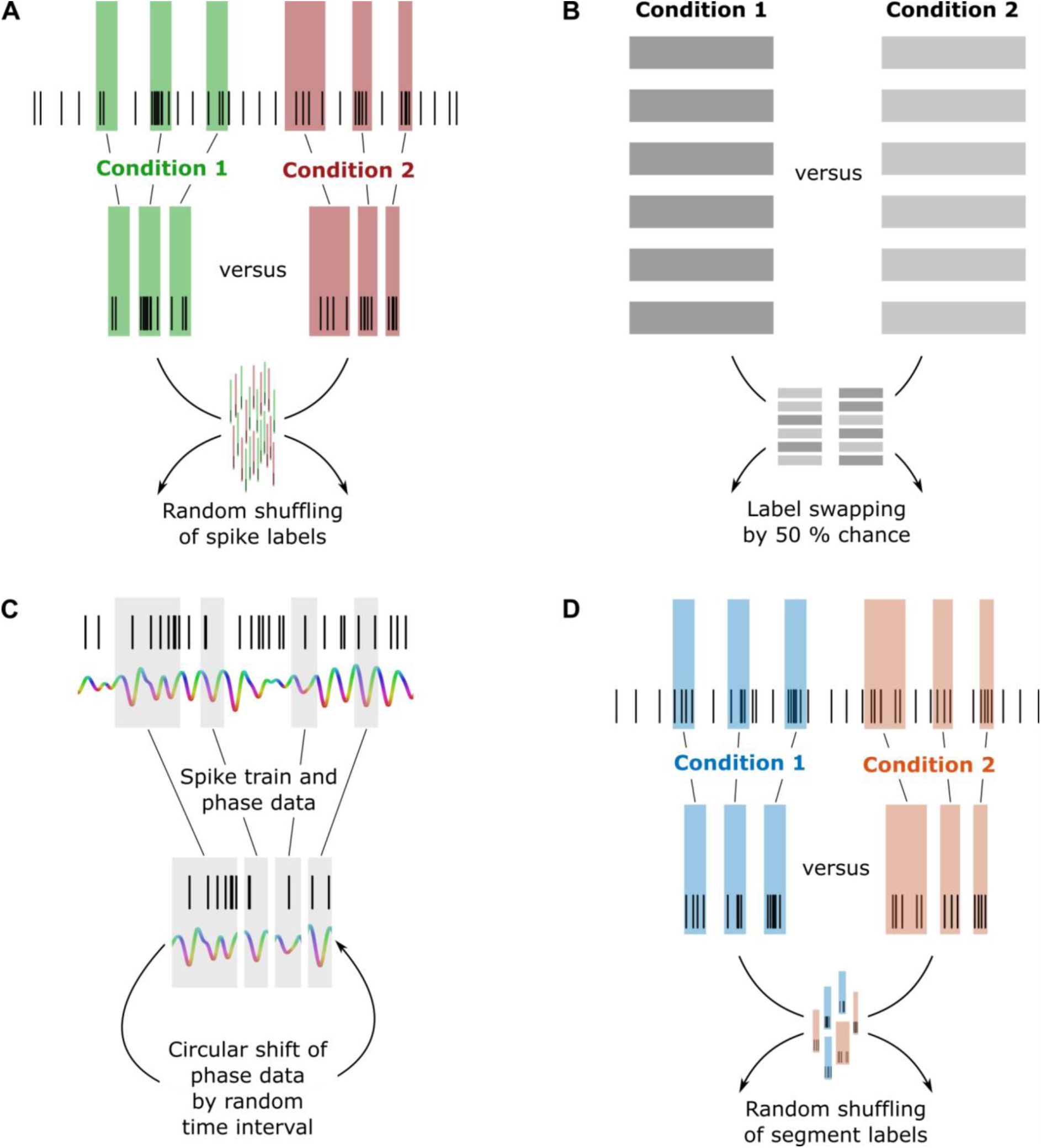
Surrogate tests. Since spike counts have an influence on several tests for assessing phase locking strength, we implemented permutation tests to account for it. (**A**) Permutation procedure for shuffling spike labels across conditions. We randomly rearranged the spike labels to create surrogate datasets. This procedure was used to maintain the original number of spikes for both conditions, for instance when comparing phase-locking strength between two conditions. (**B**) Permutation procedure with label swapping. For each pair, the labels were randomly swapped or maintained to create surrogate data. The original pairing is kept, only the assignment to the conditions is permuted. This procedure was used to compare paired data. (**C**) Permutation procedure with circular shift. Concatenated phase data was circularly shifted by random time intervals relative to the spike train to create surrogate datasets. This procedure was used to preserve the original inter-spike intervals and the basic structure of the phase data, for instance when testing the significance of a neuron’s phase locking. (**D**) Permutation procedure for shuffling segment labels across conditions. We randomly rearranged the segment labels to create surrogate datasets. This procedure was used to maintain the original number and timing of spikes within segments, for instance when comparing preferred theta phases during encoding and retrieval.

**Figure S4.**
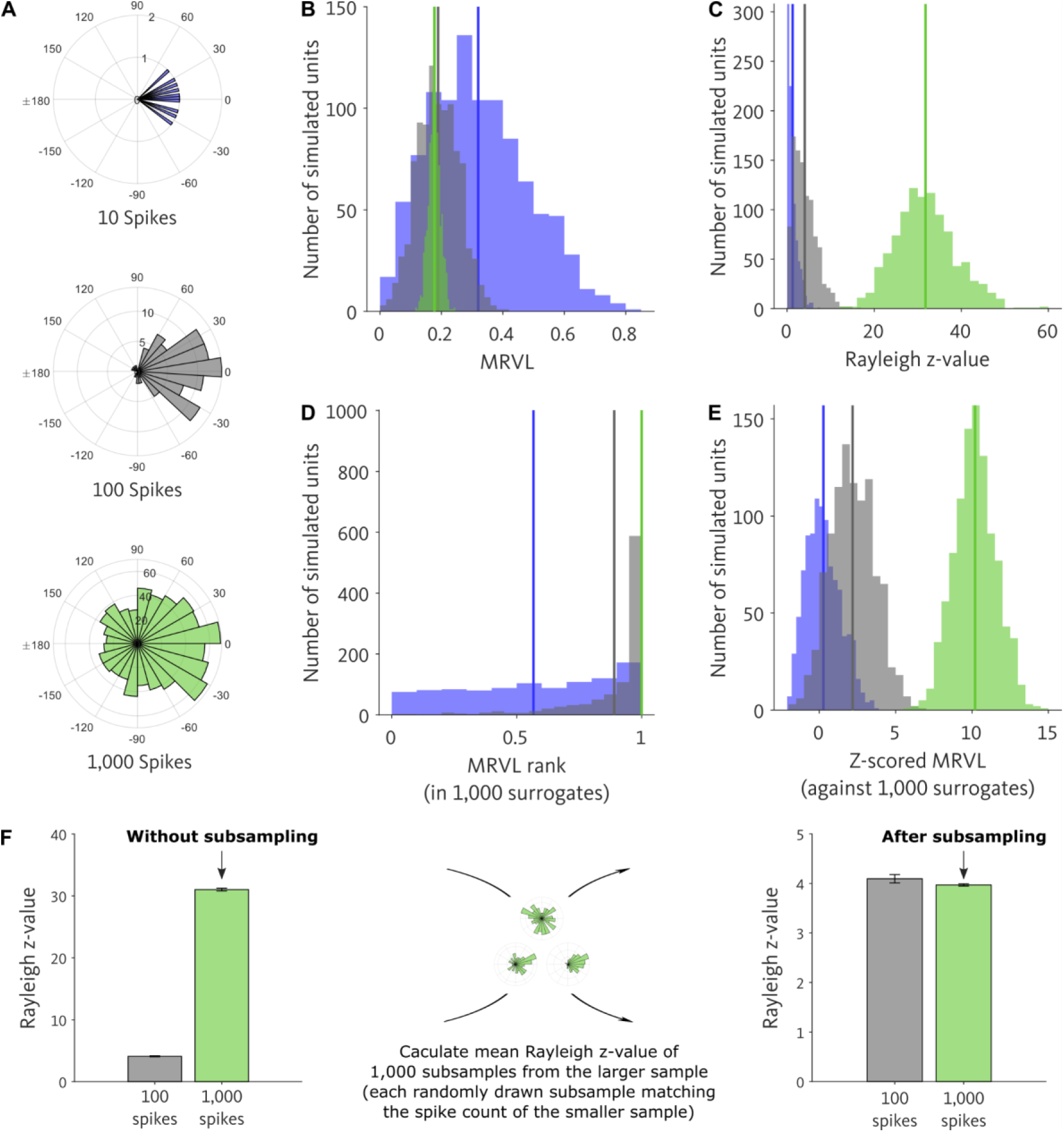
Spike counts influence test statistics for phase-locking strength. To better understand test statistics for assessing phase-locking strength, we simulated 1,000 units with phase locking and applied different test statistics on these simulated data. For each unit, we randomly sampled 250 phase angles from a Von Mises distribution centered around 0°. We added 750 completely random phase angles, resulting in a total of 1,000 phase angles with a preference towards 0°. We considered these simulated phase angles comparable to the phase angles of phase-locking units. (**A**) For each unit, we estimated phase locking-strengths on three different spike counts by separately analyzing 10 (blue), 100 (gray), and all (green) of the 1,000 phase angles. Examples for one unit are shown. (**B**) We calculated the distribution of mean resultant vector lengths (MRVLs) across units separately for the 10-spikes, 100-spikes and 1000-spikes conditions. Mean MRVLs (vertical lines) decreased with increasing numbers of spikes. (**C**) We calculated the distributions of Rayleigh *z*-values. Mean Rayleigh *z*-values (vertical lines) increased with increasing numbers of spikes. (**D**) We ranked each unit’s MRVL in a distribution of MRVLs, which were calculated from 1,000 completely random phase angle distributions with a matching spike count. Mean ranks (vertical lines) increased with increasing numbers of spikes. For 1,000 spikes, the MRVL rank was 1 for all units. (**E**) We *z*-scored each unit’s MRVL against the MRVLs from 1,000 completely random phase angle distributions with a matching spike count. Mean *z*-scored MRVLs (vertical lines) increased with increasing numbers of spikes. (**B**–**E**) In summary, all four metrics were influenced by the number of spikes. In the end, we decided to use the Rayleigh *z*-value in our analyses. Importantly, whenever we compared data from conditions with different spike counts, we controlled for the number of spikes in order to preclude that the results were simply driven by different spike counts. (**F**) To visualize our comparisons of phase-locking strength between conditions with different spike counts, we implemented a subsampling procedure. To this end, we drew random subsamples from the larger sample with each subsample matching the spike count of the smaller sample. We then calculated the mean Rayleigh *z*-value across these subsamples.

**Figure S5.**
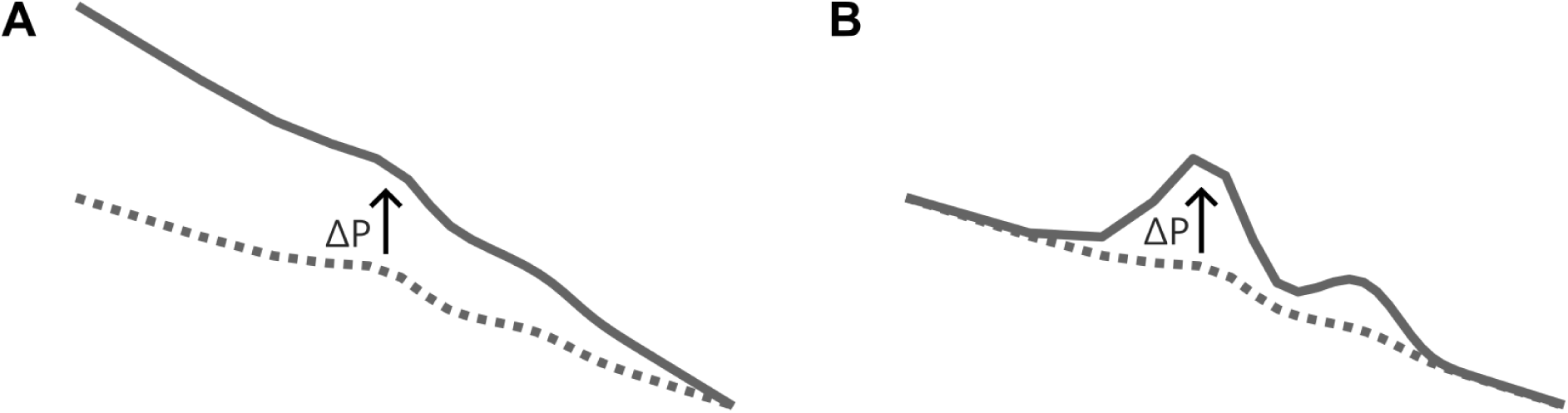
Two different causes of increases in absolute power. (**A**) Absolute power increases (ΔP > 0; black arrow) because of an increased aperiodic slope and an increased y-axis offset, indicated by the upward shift from the dotted reference line to the solid line. There is no increase in narrowband (oscillatory) power. (**B**) Absolute power increases (ΔP > 0) because of increased narrowband (oscillatory) power, indicated by the more pronounced wave above the dotted reference line. Adapted from previous studies (*6*, *55*).

**Figure S6.**
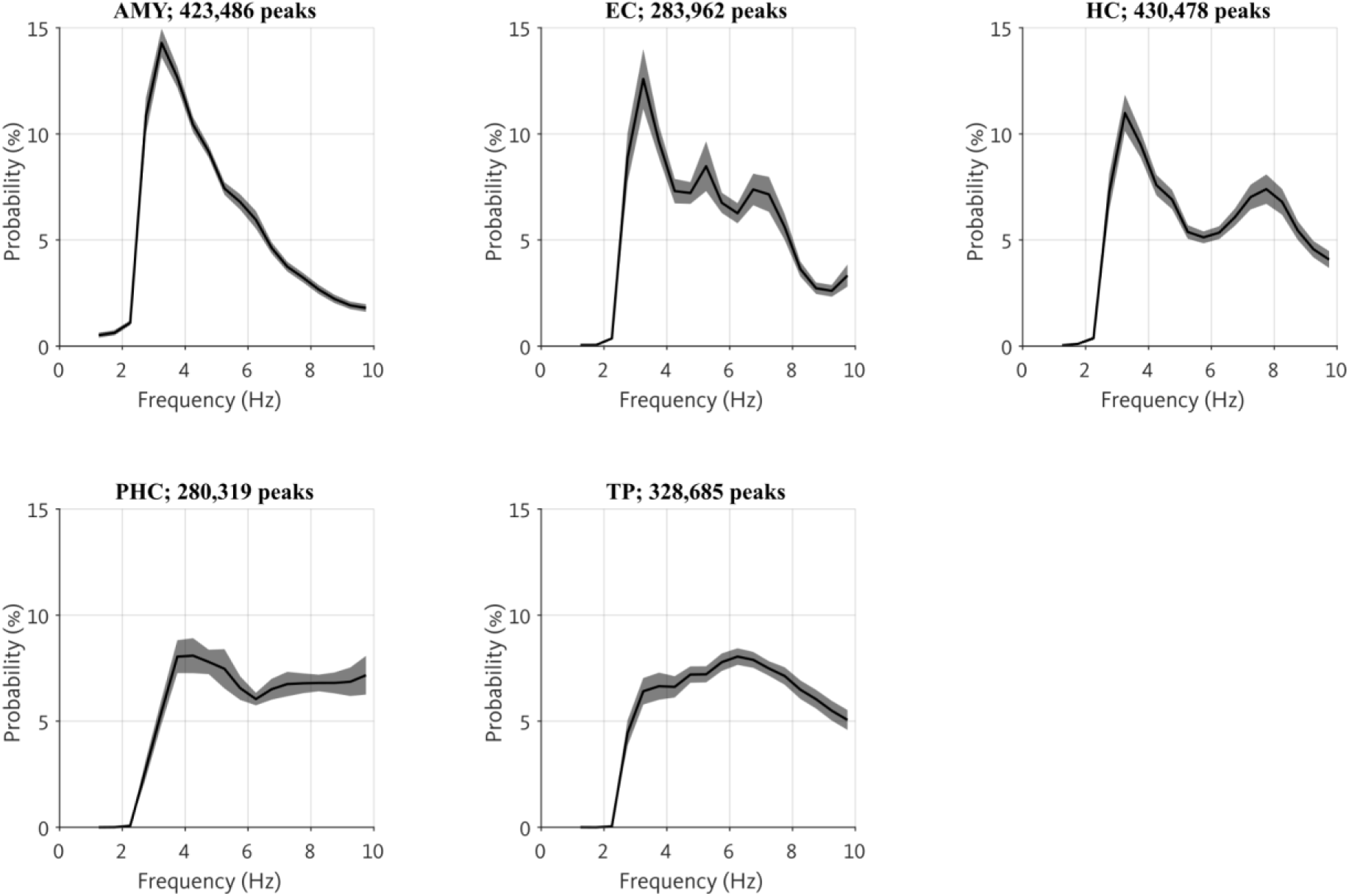
Distributions of theta peak frequencies for different brain regions. Plots show probability distributions of center frequencies of detected theta peaks across microwires, identified using SPRiNT (*56*). Gray shaded areas, standard error of the mean across microwires. AMY, amygdala; EC, entorhinal cortex; HC, hippocampus; PHC, parahippocampal cortex; TP, temporal pole.

**Figure S7.**
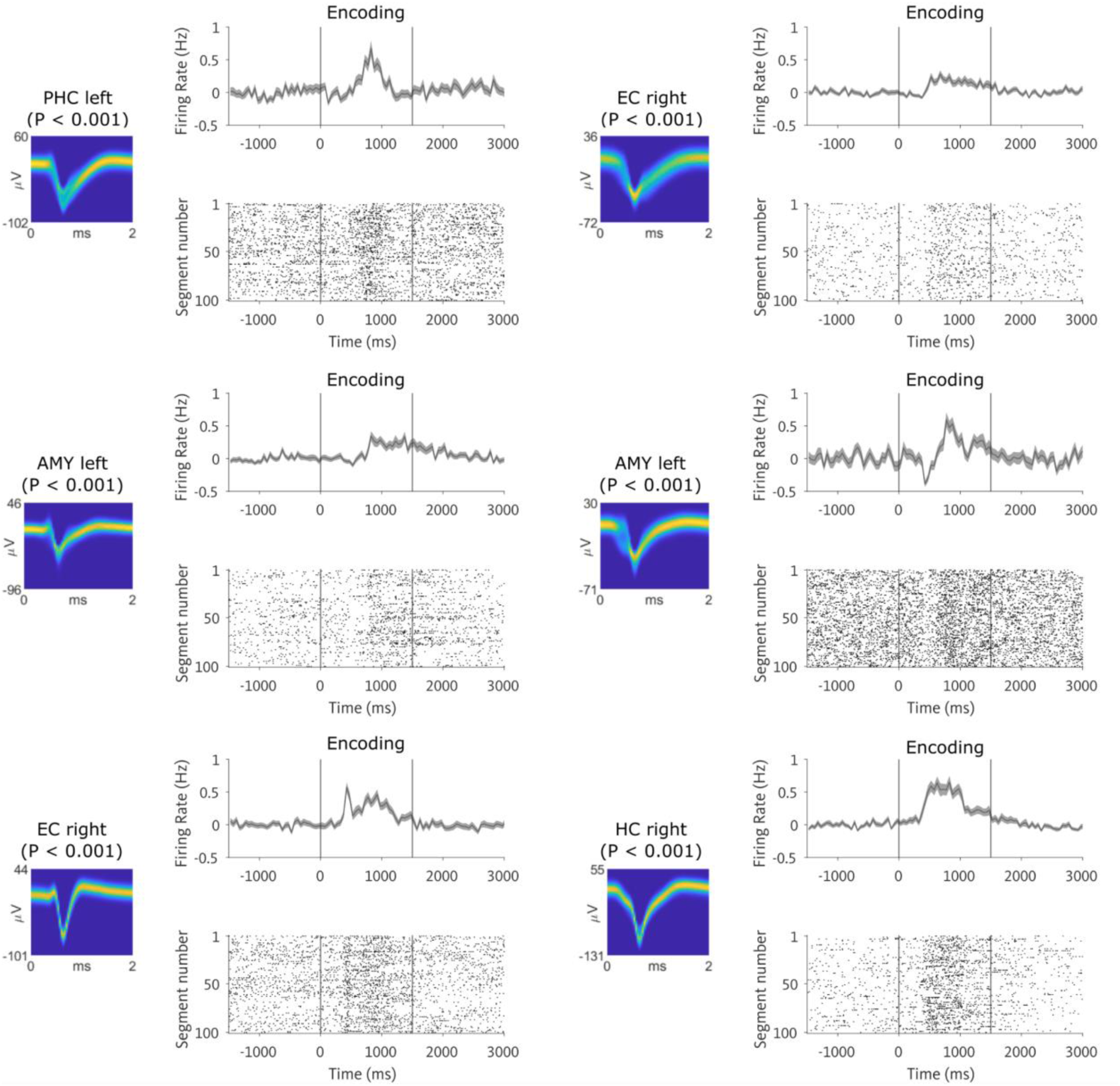
Examples of object-responsive units. We identified object-responsive units as those units that exhibited a significant increase in firing during the encoding period of the task (in comparison to baseline periods). This figure shows six example object-responsive units. For each unit, the top plot shows the mean firing rate across encoding segments. Gray shaded area indicates the standard error of the mean. Vertical dotted lines indicate the start and end time of the encoding period. Bottom raster plots show spiking activity during each encoding segment. Spike waveforms are shown as density plots on the left. AMY, amygdala; EC, entorhinal cortex; HC, hippocampus; PHC, parahippocampal cortex.

**Figure S8.**
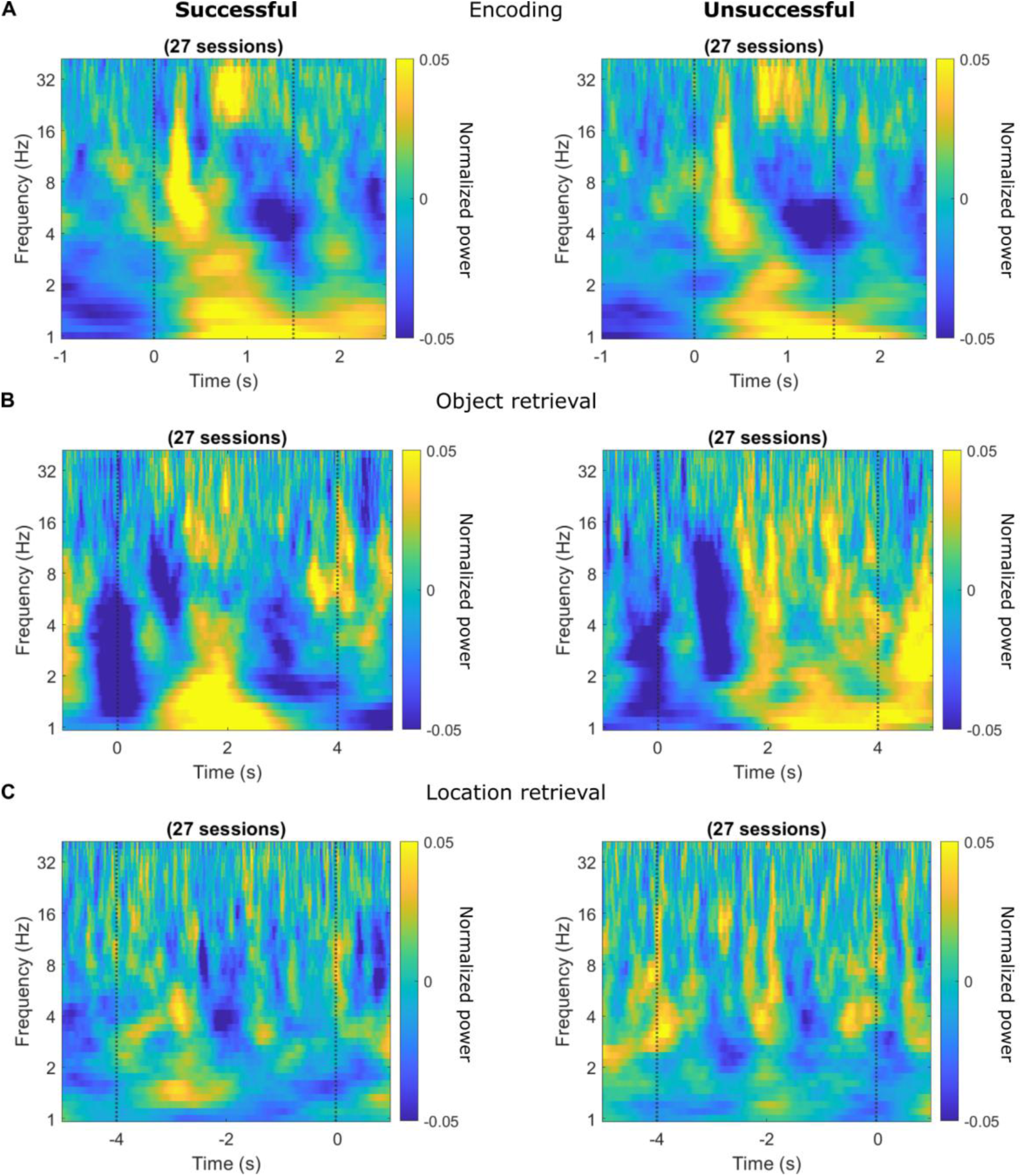
Time–frequency-resolved power during encoding and retrieval. Plots show the grand mean normalized power in the frequency range of 1 to 40 Hz during successful and unsuccessful encoding (**A**), object recall (**B**), and location recall (**C**). Dotted vertical lines indicate the start and end time of the segments. For encoding and object recall, time = 0 s indicates the start of the segment. For location recall, time = 0 s is referenced to the segment end. We included power spectrograms of all microwires with at least one unit in the analysis. We computed spectral power in time bins of 10 ms during encoding segments (1.5 s duration), object retrieval segments (4 s duration), and the last 4 s of the location retrieval segments including 5 s buffer periods before and after to avoid edge artifacts. To obtain the power spectrogram of each microwire, we calculated a continuous Morlet wavelet transform (number of cycles, 5) at 40 logarithmically spaced frequencies between 1 and 40 Hz. The resulting power values of each microwire were log-transformed and then normalized separately for each frequency and separately for encoding, object recall, and location recall (including the segments and their buffer periods). For each session, we calculated the mean power spectrum across all segments and then across all microwires with at least one unit (from various brain regions). In a last step, we calculated the grand mean across all sessions. We did not observe any significant power differences between successful and unsuccessful segments (two-sided cluster-based permutation tests, 1001 permutations: encoding (0–1.5 s), largest positive cluster, *t*(26) = 401.180, *P* = 0.120, largest negative cluster, *t*(26) = -27.238, *P* = 0.842; object recall (0–4 s), largest positive cluster, *t*(26) = 1104.220, *P* = 0.067, largest negative cluster, *t*(26) = -911.752, *P* = 0.090; location recall (-4–0 s), largest positive cluster, *t*(26) = 96.496, *P* = 0.685, largest negative cluster, *t*(26) = -686.795, *P* = 0.162).

**Figure S9.**
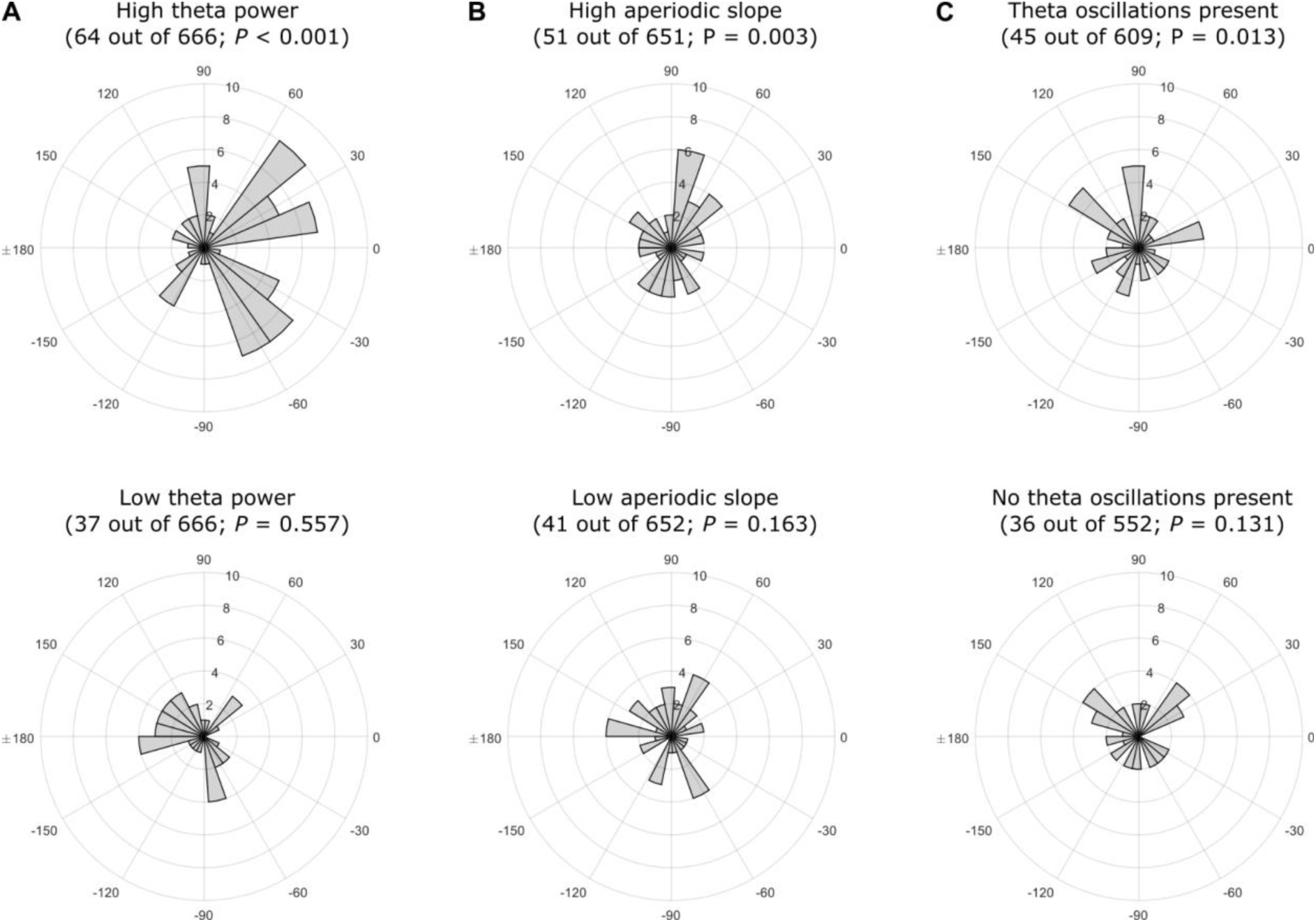
Encoding–retrieval phase shifts as a function of theta power, aperiodic slope, and theta oscillations. Polar histograms show angular theta-phase differences of neurons with significant theta-phase shifts between encoding and retrieval. For different conditions, the number of phase-shifting units and the *P*-values of a binomial test are displayed. (**A**) Phase shifts for high and low theta power. (**B**) Phase shifts for high and low aperiodic slope. (**C**) Phase shifts in the presence and absence of theta oscillations.

**Table S1.** Participant information.

| Participant index | Session index | Number of trials | Number of microwires | Number of units |
| --- | --- | --- | --- | --- |
| 1 | 1 | 21 | 32 | 19 |
| 1 | 2 | 39 | 32 | 18 |
| 2 | 1 | 32 | 64 | 64 |
| 2 | 2 | 32 | 64 | 63 |
| 3 | 1 | 40 | 32 | 50 |
| 3 | 2 | 28 | 32 | 55 |
| 4 | 1 | 40 | 24 | 28 |
| 4 | 2 | 40 | 24 | 12 |
| 5 | 1 | 40 | 48 | 48 |
| 5 | 2 | 40 | 48 | 40 |
| 6 | 1 | 40 | 48 | 33 |
| 6 | 2 | 40 | 48 | 23 |
| 7 | 1 | 24 | 48 | 52 |
| 8 | 1 | 32 | 48 | 42 |
| 8 | 2 | 24 | 48 | 36 |
| 9 | 1 | 40 | 48 | 13 |
| 10 | 1 | 40 | 48 | 42 |
| 10 | 2 | 40 | 48 | 37 |
| 11 | 1 | 40 | 48 | 48 |
| 12 | 1 | 40 | 48 | 56 |
| 13 | 1 | 40 | 16 | 19 |
| 14 | 1 | 40 | 48 | 48 |
| 15 | 1 | 40 | 32 | 24 |
| 16 | 1 | 35 | 32 | 45 |
| 17 | 1 | 40 | 48 | 62 |
| 18 | 1 | 40 | 32 | 28 |
| 18 | 2 | 40 | 32 | 20 |

**Table S2.** Fixed effects estimates for the effect of brain regions on Rayleigh *z*-values in a linear mixed-effects model. Model formula: *z*-value ∼ 1 + regions + (1|sessions) + (1|periods). Note that the effect was driven by increased Rayleigh *z*-values in the parahippocampal cortex.

| **Predictor** | **Degrees of freedom** | **t-value** | ***P*-value** |
| --- | --- | --- | --- |
| **AMY versus EC** | 1894 | -2.457 | 0.014 |
| **AMY versus HC** | 1894 | 0.1 | 0.92 |
| **AMY versus PHC** | 1894 | -7.4 | <0.001 |
| **AMY versus TP** | 1894 | -1.032 | 0.302 |
| **EC versus HC** | 1894 | 2.249 | 0.025 |
| **EC versus PHC** | 1894 | -4.829 | <0.001 |
| **EC versus TP** | 1894 | 1.064 | 0.287 |
| **HC versus PHC** | 1894 | -6.863 | <0.001 |
| **HC versus TP** | 1894 | -0.98 | 0.327 |
| **PHC versus TP** | 1894 | 5.452 | <0.001 |

**Table S3.** Fixed effects estimates for the effect of brain regions on mean log-transformed power assigned to spikes in a linear mixed-effects model. Model formula: power ∼ 1 + regions + (1|sessions) + (1|periods). Note that the effects were driven by increased theta power in the hippocampus.

| **Predictor** | **Degrees of freedom** | **t-value** | **P-value** |
| --- | --- | --- | --- |
| **AMY versus EC** | 1894 | -13.738 | <0.001 |
| **AMY versus HC** | 1894 | -26.237 | <0.001 |
| **AMY versus PHC** | 1894 | -9.253 | <0.001 |
| **AMY versus TP** | 1894 | -17.29 | <0.001 |
| **EC versus HC** | 1894 | -11.014 | <0.001 |
| **EC versus PHC** | 1894 | 2.572 | 0.010 |
| **EC versus TP** | 1894 | -3.789 | <0.001 |
| **HC versus PHC** | 1894 | 12.773 | <0.001 |
| **HC versus TP** | 1894 | 6.32 | <0.001 |
| **PHC versus TP** | 1894 | -5.938 | <0.001 |

## Notes

https://github.com/NeuroGuth/GuthPhaseLocking2024

### Competing Interest Statement

The authors have declared no competing interest.

## References

1. G. Buzsáki, C. A. Anastassiou, C. Koch, The origin of extracellular fields and currents — EEG, ECoG, LFP and spikes. Nat. Rev. Neurosci. 13, 407–420 (2012).

2. S. Katzner, I. Nauhaus, A. Benucci, V. Bonin, D. L. Ringach, M. Carandini, Local Origin of Field Potentials in Visual Cortex. Neuron 61, 35–41 (2009).

3. S. E. Qasim, L. Kunz, “How Is Single-Neuron Activity Related to LFP Oscillations?” in Intracranial EEG: A Guide for Cognitive Neuroscientists, N. Axmacher, Ed. (Springer International Publishing, Cham, 2023; 10.1007/978-3-031-20910-9_44) *Studies in Neuroscience, Psychology and Behavioral Economics*, pp. 703–718.

4. H. G. Rey, C. Pedreira, R. Quian Quiroga, Past, present and future of spike sorting techniques. Brain Res. Bull. 119, 106–117 (2015).

5. G. Buzsáki, A. Draguhn, Neuronal Oscillations in Cortical Networks. Science 304, 1926–1929 (2004).

6. N. A. Herweg, E. A. Solomon, M. J. Kahana, Theta Oscillations in Human Memory. Trends Cogn. Sci. 24, 208–227 (2020).

7. G. Buzsáki, Theta Oscillations in the Hippocampus. Neuron 33, 325–340 (2002).

8. J. Jacobs, Hippocampal theta oscillations are slower in humans than in rodents: implications for models of spatial navigation and memory. Philos. Trans. R. Soc. B Biol. Sci. 369, 20130304 (2014).

9. L. Kunz, S. Maidenbaum, D. Chen, L. Wang, J. Jacobs, N. Axmacher, Mesoscopic Neural Representations in Spatial Navigation. Trends Cogn. Sci. 23, 615–630 (2019).

10. A. Nuñez, W. Buño, The Theta Rhythm of the Hippocampus: From Neuronal and Circuit Mechanisms to Behavior. Front. Cell. Neurosci. 15, 649262 (2021).

11. M. J. Bezaire, I. Raikov, K. Burk, D. Vyas, I. Soltesz, Interneuronal mechanisms of hippocampal theta oscillations in a full-scale model of the rodent CA1 circuit. eLife 5, e18566 (2016).

12. G. Buzsáki, Theta rhythm of navigation: Link between path integration and landmark navigation, episodic and semantic memory. Hippocampus 15, 827–840 (2005).

13. J. O’Keefe, M. L. Recce, Phase relationship between hippocampal place units and the EEG theta rhythm. Hippocampus 3, 317–330 (1993).

14. L. L. Colgin, Rhythms of the hippocampal network. Nat. Rev. Neurosci. 17, 239–249 (2016).

15. C. H. Vanderwolf, Hippocampal electrical activity and voluntary movement in the rat. Electroencephalogr. Clin. Neurophysiol. 26, 407–418 (1969).

16. A. J. Watrous, D. J. Lee, A. Izadi, G. G. Gurkoff, K. Shahlaie, A. D. Ekstrom, A comparative study of human and rat hippocampal low-frequency oscillations during spatial navigation. Hippocampus 23, 656–661 (2013).

17. J. Winson, Loss of Hippocampal Theta Rhythm Results in Spatial Memory Deficit in the Rat. Science 201, 160–163 (1978).

18. F. Stella, A. Treves, Associative Memory Storage and Retrieval: Involvement of Theta Oscillations in Hippocampal Information Processing. Neural Plast. 2011, 1–15 (2011).

19. K. Mizuseki, A. Sirota, E. Pastalkova, G. Buzsáki, Theta Oscillations Provide Temporal Windows for Local Circuit Computation in the Entorhinal-Hippocampal Loop. Neuron 64, 267– 280 (2009).

20. T. Gedankien, R. J. Tan, S. E. Qasim, H. Moore, D. McDonagh, J. Jacobs, B. Lega, Acetylcholine modulates the temporal dynamics of human theta oscillations during memory. Nat. Commun. 14, 5283 (2023).

21. T. Korotkova, A. Ponomarenko, C. K. Monaghan, S. L. Poulter, F. Cacucci, T. Wills, M. E. Hasselmo, C. Lever, Reconciling the different faces of hippocampal theta: The role of theta oscillations in cognitive, emotional and innate behaviors. Neurosci. Biobehav. Rev. 85, 65–80 (2018).

22. A. J. Watrous, J. Miller, S. E. Qasim, I. Fried, J. Jacobs, Phase-tuned neuronal firing encodes human contextual representations for navigational goals. eLife 7, e32554 (2018).

23. M. J. Kahana, R. Sekuler, J. B. Caplan, M. Kirschen, J. R. Madsen, Human theta oscillations exhibit task dependence during virtual maze navigation. Nature 399, 781–784 (1999).

24. V. D. Bohbot, M. S. Copara, J. Gotman, A. D. Ekstrom, Low-frequency theta oscillations in the human hippocampus during real-world and virtual navigation. Nat. Commun. 8, 14415 (2017).

25. Z. M. Aghajan, P. Schuette, T. A. Fields, M. E. Tran, S. M. Siddiqui, N. R. Hasulak, T. K. Tcheng, D. Eliashiv, E. A. Mankin, J. Stern, I. Fried, N. Suthana, Theta Oscillations in the Human Medial Temporal Lobe during Real-World Ambulatory Movement. Curr. Biol. 27, 3743–3751.e3 (2017).

26. A. Goyal, J. Miller, S. E. Qasim, A. J. Watrous, H. Zhang, J. M. Stein, C. S. Inman, R. E. Gross, J. T. Willie, B. Lega, J.-J. Lin, A. Sharan, C. Wu, M. R. Sperling, S. A. Sheth, G. M. McKhann, E. H. Smith, C. Schevon, J. Jacobs, Functionally distinct high and low theta oscillations in the human hippocampus. Nat. Commun. 11, 2469 (2020).

27. M. Stangl, U. Topalovic, C. S. Inman, S. Hiller, D. Villaroman, Z. M. Aghajan, L. Christov-Moore, N. R. Hasulak, V. R. Rao, C. H. Halpern, D. Eliashiv, I. Fried, N. Suthana, Boundary-anchored neural mechanisms of location-encoding for self and others. Nature, 1–6 (2020).

28. B. C. Lega, J. Jacobs, M. Kahana, Human hippocampal theta oscillations and the formation of episodic memories. Hippocampus 22, 748–761 (2012).

29. A. J. Watrous, I. Fried, A. D. Ekstrom, Behavioral correlates of human hippocampal delta and theta oscillations during navigation. J. Neurophysiol. 105, 1747–1755 (2011).

30. L. Kunz, L. Wang, D. Lachner-Piza, H. Zhang, A. Brandt, M. Dümpelmann, P. C. Reinacher, V. A. Coenen, D. Chen, W.-X. Wang, W. Zhou, S. Liang, P. Grewe, C. G. Bien, A. Bierbrauer, T. Navarro Schröder, A. Schulze-Bonhage, N. Axmacher, Hippocampal theta phases organize the reactivation of large-scale electrophysiological representations during goal-directed navigation. Sci. Adv. 5, eaav8192 (2019).

31. A. Navas-Olive, M. Valero, T. Jurado-Parras, A. de Salas-Quiroga, R. G. Averkin, G. Gambino, E. Cid, L. M. de la Prida, Multimodal determinants of phase-locked dynamics across deep-superficial hippocampal sublayers during theta oscillations. Nat. Commun. 11, 2217 (2020).

32. A. G. Siapas, E. V. Lubenov, M. A. Wilson, Prefrontal Phase Locking to Hippocampal Theta Oscillations. Neuron 46, 141–151 (2005).

33. A. Sirota, S. Montgomery, S. Fujisawa, Y. Isomura, M. Zugaro, G. Buzsáki, Entrainment of Neocortical Neurons and Gamma Oscillations by the Hippocampal Theta Rhythm. Neuron 60, 683–697 (2008).

34. J. Jacobs, M. J. Kahana, A. D. Ekstrom, I. Fried, Brain Oscillations Control Timing of Single-Neuron Activity in Humans. J. Neurosci. 27, 3839–3844 (2007).

35. U. Rutishauser, I. B. Ross, A. N. Mamelak, E. M. Schuman, Human memory strength is predicted by theta-frequency phase-locking of single neurons. Nature 464, 903–907 (2010).

36. D. R. Schonhaut, A. M. Rao, A. G. Ramayya, E. A. Solomon, N. A. Herweg, I. Fried, M. J. Kahana, MTL neurons phase-lock to human hippocampal theta. eLife 13, e85753 (2024).

37. T. Hafting, M. Fyhn, T. Bonnevie, M.-B. Moser, E. I. Moser, Hippocampus-independent phase precession in entorhinal grid cells. Nature 453, 1248–1252 (2008).

38. W. E. Skaggs, B. L. McNaughton, M. A. Wilson, C. A. Barnes, Theta phase precession in hippocampal neuronal populations and the compression of temporal sequences. Hippocampus 6, 149–172 (1996).

39. S. E. Qasim, I. Fried, J. Jacobs, Phase precession in the human hippocampus and entorhinal cortex. Cell 184, 3242–3255.e10 (2021).

40. E. T. Reifenstein, C. L. Ebbesen, Q. Tang, M. Brecht, S. Schreiber, R. Kempter, Cell-Type Specific Phase Precession in Layer II of the Medial Entorhinal Cortex. J. Neurosci. 36, 2283– 2288 (2016).

41. L. Reddy, M. W. Self, B. Zoefel, M. Poncet, J. K. Possel, J. C. Peters, J. C. Baayen, S. Idema, R. VanRullen, P. R. Roelfsema, Theta-phase dependent neuronal coding during sequence learning in human single neurons. Nat. Commun. 12, 4839 (2021).

42. D. Santos-Pata, C. Barry, H. F. Ólafsdóttir, Theta-band phase locking during encoding leads to coordinated entorhinal-hippocampal replay. Curr. Biol. 33, 4570–4581.e5 (2023).

43. T. Shuman, D. Aharoni, D. J. Cai, C. R. Lee, S. Chavlis, L. Page-Harley, L. M. Vetere, Y. Feng, C. Y. Yang, I. Mollinedo-Gajate, L. Chen, Z. T. Pennington, J. Taxidis, S. E. Flores, K. Cheng, M. Javaherian, C. C. Kaba, N. Rao, M. La-Vu, I. Pandi, M. Shtrahman, K. I. Bakhurin, S. C. Masmanidis, B. S. Khakh, P. Poirazi, A. J. Silva, P. Golshani, Breakdown of spatial coding and interneuron synchronization in epileptic mice. Nat. Neurosci. 23, 229–238 (2020).

44. O. Kornienko, P. Latuske, M. Bassler, L. Kohler, K. Allen, Non-rhythmic head-direction cells in the parahippocampal region are not constrained by attractor network dynamics. eLife 7, e35949 (2018).

45. E. L. Newman, M. E. Hasselmo, Grid cell firing properties vary as a function of theta phase locking preferences in the rat medial entorhinal cortex. Front. Syst. Neurosci. 8 (2014).

46. O. Jensen, Reading the hippocampal code by theta phase-locking. Trends Cogn. Sci. 9, 551–553 (2005).

47. M. E. Hasselmo, C. Bodelón, B. P. Wyble, A Proposed Function for Hippocampal Theta Rhythm: Separate Phases of Encoding and Retrieval Enhance Reversal of Prior Learning. Neural Comput. 14, 793–817 (2002).

48. M. E. Hasselmo, C. E. Stern, Theta rhythm and the encoding and retrieval of space and time. NeuroImage 85, 656–666 (2014).

49. H. B. Yoo, G. Umbach, B. Lega, Neurons in the human medial temporal lobe track multiple temporal contexts during episodic memory processing. NeuroImage 245, 118689 (2021).

50. L. Kunz, A. Brandt, P. C. Reinacher, B. P. Staresina, E. T. Reifenstein, C. T. Weidemann, N. A. Herweg, A. Patel, M. Tsitsiklis, R. Kempter, M. J. Kahana, A. Schulze-Bonhage, J. Jacobs, A neural code for egocentric spatial maps in the human medial temporal lobe. Neuron 109, 2781–2796.e10 (2021).

51. M. Tsitsiklis, J. Miller, S. E. Qasim, C. S. Inman, R. E. Gross, J. T. Willie, E. H. Smith, S. A. Sheth, C. A. Schevon, M. R. Sperling, A. Sharan, J. M. Stein, J. Jacobs, Single-Neuron Representations of Spatial Targets in Humans. Curr. Biol. 30, 245–253.e4 (2020).

52. J. Miller, A. J. Watrous, M. Tsitsiklis, S. A. Lee, S. A. Sheth, C. A. Schevon, E. H. Smith, M. R. Sperling, A. Sharan, A. A. Asadi-Pooya, G. A. Worrell, S. Meisenhelter, C. S. Inman, K. A. Davis, B. Lega, P. A. Wanda, S. R. Das, J. M. Stein, R. Gorniak, J. Jacobs, Lateralized hippocampal oscillations underlie distinct aspects of human spatial memory and navigation. Nat. Commun. 9, 2423 (2018).

53. S. Maidenbaum, J. Miller, J. M. Stein, J. Jacobs, Grid-like hexadirectional modulation of human entorhinal theta oscillations. Proc. Natl. Acad. Sci. 115, 10798–10803 (2018).

54. Z. W. Davis, L. Muller, J. Martinez-Trujillo, T. Sejnowski, J. H. Reynolds, Spontaneous travelling cortical waves gate perception in behaving primates. Nature 587, 432–436 (2020).

55. T. Donoghue, M. Haller, E. J. Peterson, P. Varma, P. Sebastian, R. Gao, T. Noto, A. H. Lara, J. D. Wallis, R. T. Knight, A. Shestyuk, B. Voytek, Parameterizing neural power spectra into periodic and aperiodic components. Nat. Neurosci. 23, 1655–1665 (2020).

56. L. E. Wilson, J. Da Silva Castanheira, S. Baillet, Time-resolved parameterization of aperiodic and periodic brain activity. eLife 11, e77348 (2022).

57. I. Fried, C. L. Wilson, N. T. Maidment, J. Engel, E. Behnke, T. A. Fields, K. A. Macdonald, J. W. Morrow, L. Ackerson, Cerebral microdialysis combined with single-neuron and electroencephalographic recording in neurosurgical patients: Technical note. J. Neurosurg. 91, 697–705 (1999).

58. I. Fried, U. Rutishauser, M. Cerf, G. Kreiman, Single Neuron Studies of the Human Brain: Probing Cognition (MIT Press, Cambridge, MA, USA, 2014).

59. U. Rutishauser, L. Reddy, F. Mormann, J. Sarnthein, The Architecture of Human Memory: Insights from Human Single-Neuron Recordings. J. Neurosci. 41, 883–890 (2021).

60. J. Jacobs, M. J. Kahana, Direct brain recordings fuel advances in cognitive electrophysiology. Trends Cogn. Sci. 14, 162–171 (2010).

61. R. Quiroga, Plugging in to Human Memory: Advantages, Challenges, and Insights from Human Single-Neuron Recordings. Cell 179, 1015–1032 (2019).

62. R. Gao, E. J. Peterson, B. Voytek, Inferring synaptic excitation/inhibition balance from field potentials. NeuroImage 158, 70–78 (2017).

63. D. Bush, J. A. Bisby, C. M. Bird, S. Gollwitzer, R. Rodionov, B. Diehl, A. W. McEvoy, M. C. Walker, N. Burgess, Human hippocampal theta power indicates movement onset and distance travelled. Proc. Natl. Acad. Sci. 114, 12297–12302 (2017).

64. V. Douchamps, A. Jeewajee, P. Blundell, N. Burgess, C. Lever, Evidence for Encoding versus Retrieval Scheduling in the Hippocampus by Theta Phase and Acetylcholine. J. Neurosci. 33, 8689–8704 (2013).

65. L. L. Colgin, Oscillations and hippocampal–prefrontal synchrony. Curr. Opin. Neurobiol. 21, 467–474 (2011).

66. J. E. Lisman, O. Jensen, The Theta-Gamma Neural Code. Neuron 77, 1002–1016 (2013).

67. G. Buzsáki, Neural Syntax: Cell Assemblies, Synapsembles, and Readers. Neuron 68, 362–385 (2010).

68. M. E. Hasselmo, What is the function of hippocampal theta rhythm?—Linking behavioral data to phasic properties of field potential and unit recording data. Hippocampus 15, 936–949 (2005).

69. C. C. Canavier, Phase-resetting as a tool of information transmission. Curr. Opin. Neurobiol. 31, 206–213 (2015).

70. J. P. Kennedy, Y. Zhou, Y. Qin, S. D. Lovett, A. Sheremet, S. N. Burke, A. P. Maurer, A Direct Comparison of Theta Power and Frequency to Speed and Acceleration. J. Neurosci. 42, 4326– 4341 (2022).

71. M. Kopčanová, L. Tait, T. Donoghue, G. Stothart, L. Smith, A. A. Flores-Sandoval, P. Davila-Perez, S. Buss, M. M. Shafi, A. Pascual-Leone, P. J. Fried, C. S. Y. Benwell, Resting-state EEG signatures of Alzheimer’s disease are driven by periodic but not aperiodic changes. Neurobiol. Dis. 190, 106380 (2024).

72. E. Cesnaite, P. Steinfath, M. Jamshidi Idaji, T. Stephani, D. Kumral, S. Haufe, C. Sander, T. Hensch, U. Hegerl, S. Riedel-Heller, S. Röhr, M. L. Schroeter, A. Witte, A. Villringer, V. V. Nikulin, Alterations in rhythmic and non-rhythmic resting-state EEG activity and their link to cognition in older age. NeuroImage 268, 119810 (2023).

73. A. T. Hill, G. M. Clark, F. J. Bigelow, J. A. G. Lum, P. G. Enticott, Periodic and aperiodic neural activity displays age-dependent changes across early-to-middle childhood. Dev. Cogn. Neurosci. 54, 101076 (2022).

74. S. Kota, M. D. Rugg, B. C. Lega, Hippocampal Theta Oscillations Support Successful Associative Memory Formation. J. Neurosci. 40, 9507–9518 (2020).

75. T. A. Whitten, A. M. Hughes, C. T. Dickson, J. B. Caplan, A better oscillation detection method robustly extracts EEG rhythms across brain state changes: The human alpha rhythm as a test case. NeuroImage 54, 860–874 (2011).

76. L. Kunz, B. P. Staresina, P. C. Reinacher, A. Brandt, T. A. Guth, A. Schulze-Bonhage, J. Jacobs, Ripple-locked coactivity of stimulus-specific neurons and human associative memory. Nat. Neurosci. 27, 587–599 (2024).

77. F. J. Chaure, H. G. Rey, R. Quian Quiroga, A novel and fully automatic spike-sorting implementation with variable number of features. J. Neurophysiol. 120, 1859–1871 (2018).

78. T. A. Guth, L. Kunz, A. Brandt, M. Dümpelmann, K. A. Klotz, P. C. Reinacher, A. Schulze-Bonhage, J. Jacobs, J. Schönberger, Interictal spikes with and without high-frequency oscillation have different single-neuron correlates. Brain 144, 3078–3088 (2021).

79. R. Q. Quiroga, R. Mukamel, E. A. Isham, R. Malach, I. Fried, Human single-neuron responses at the threshold of conscious recognition. Proc. Natl. Acad. Sci. 105, 3599–3604 (2008).

80. Z. W. Davis, L. Muller, J. H. Reynolds, Spontaneous Spiking Is Governed by Broadband Fluctuations. J. Neurosci. 42, 5159–5172 (2022).

81. P. Fries, P. R. Roelfsema, A. K. Engel, P. König, W. Singer, Synchronization of oscillatory responses in visual cortex correlates with perception in interocular rivalry. Proc. Natl. Acad. Sci. 94, 12699–12704 (1997).

82. P. P. Mitra, B. Pesaran, Analysis of Dynamic Brain Imaging Data. Biophys. J. 76, 691–708 (1999).

83. R. Oostenveld, P. Fries, E. Maris, J.-M. Schoffelen, FieldTrip: Open Source Software for Advanced Analysis of MEG, EEG, and Invasive Electrophysiological Data. Comput. Intell. Neurosci. 2011, e156869 (2010).

84. P. Berens, CircStat: A MATLAB Toolbox for Circular Statistics. J. Stat. Softw. 31, 1–21 (2009).

